# The early macrophage response to pathogens requires dynamic regulation of the nuclear paraspeckle

**DOI:** 10.1101/2023.05.11.540384

**Authors:** Sikandar Azam, Kaitlyn S. Armijo, Chi G. Weindel, Alice Devigne, Shinichi Nakagawa, Tetsuro Hirose, Susan Carpenter, Robert O. Watson, Kristin L. Patrick

## Abstract

To ensure a robust immune response to pathogens without risking immunopathology, the kinetics and amplitude of inflammatory gene expression in macrophages needs to be exquisitely well-controlled. There is a growing appreciation for stress-responsive membraneless organelles (MLOs) regulating various steps of eukaryotic gene expression in response to extrinsic cues. Here, we implicate the nuclear paraspeckle, a highly ordered biomolecular condensate that nucleates on the *Neat1* lncRNA, in tuning innate immune gene expression in murine macrophages. In response to a variety of innate agonists, macrophage paraspeckles rapidly aggregate (0.5 h post-stimulation) and disaggregate (2h post-stimulation). Paraspeckle maintenance and aggregation require active transcription and MAPK signaling whereas paraspeckle disaggregation requires degradation of *Neat1* via the nuclear RNA exosome. Expression of a large cohort of cytokines, chemokines, and antimicrobial mediators is compromised in lipopolysaccharide-treated macrophages lacking *Neat1*, resulting in a failure to express a cohort of pro-inflammatory cytokines, chemokines, and antimicrobial mediators. Consequently, *Neat1* KO macrophages cannot control replication of *Salmonella enterica* serovar Typhimurium or vesicular stomatitis virus. These findings highlight a prominent role for MLOs in orchestrating the macrophage response to pathogens and support a model whereby dynamic assembly and disassembly of paraspeckles reprograms the nuclear RNA binding protein landscape to enable inflammatory gene expression following innate stimuli.

**SIGNIFICANCE STATEMENT:** To mount appropriate immune responses and fight infection, macrophages need to sense and respond to pathogen-associated signals with incredible precision. Membraneless organelles (MLOs) are complexes of RNAs and proteins that change in size, shape, and abundance in response to extracellular signals. We hypothesized that an MLO called the nuclear paraspeckle helps macrophages initiate and calibrate innate immune gene expression during infection. We found that paraspeckles rapidly aggregate and then dissolve in macrophages following pathogen sensing. Macrophages lacking paraspeckles cannot properly induce inflammatory genes, resulting in a failure to control replication of intracellular bacterial and viral pathogens. These data suggest that altered paraspeckle dynamics may dysregulate inflammatory gene expression in a variety of human diseases.

## INTRODUCTION

Biologists have long been interested in the functions of the membrane-bound organelles that define eukaryotic cells. More recently, membrane*less* organelles (MLO) have captured the attention of many. MLOs are biomolecular condensates that form through the process of liquid-liquid phase separation (LLPS). Through sequestration of proteins and RNAs, MLOs regulate and compartmentalize a variety of cellular processes in both the cytosol and the nucleus. The nucleus contains several MLOs, including the nucleolus, Cajal bodies, PML nuclear bodies, nuclear speckles, and nuclear paraspeckles. These nuclear condensates regulate many steps in gene expression, including transcription, pre-mRNA splicing, RNA editing, and mRNA export (1–3). A common theme of MLOs is their ability to change number, size, structure, and composition in response to cellular stress (4, 5). While phenomena related to condensate assembly have been extensively described, mechanistic links between MLO dynamics and function remain poorly understood.

One nuclear condensate with well-established links to stress responses is the nuclear paraspeckle (6–8). Discovered in HeLa cells in 2002, nuclear paraspeckles were first defined as nuclear domains enriched for paraspeckle protein 1 (PSP1) found in proximity to SC35-containing nuclear speckles (7). Paraspeckles are characterized by their spheroidal shape and distinct core and shell-like structure (7, 9, 10). These highly ordered MLOs organize on a long lncRNA called nuclear paraspeckle assembly transcript 1 (*Neat1*) (10). The *Neat1* gene encodes two isoforms, *Neat1*_*1* and *Neat1*_*2*. While both are found in paraspeckles, only *Neat1*_*2*, the long isoform (22.7 kb in humans, 21.2 kb in mice), is required for paraspeckle assembly. Although the two isoforms share the same promoter, their processing is distinct; instead of being polyadenylated, the 3’ end of *Neat1_2* is stabilized by an atypical triple helix structure (8, 11). The short *Neat1*_*1* isoform, on the other hand, is spliced and polyadenylated (1, 10).

Paraspeckles are comprised of ∼50 copies of *Neat1*_*2* and a cohort of RNA binding proteins (RBPs), several of which are required for paraspeckle assembly/maintenance. The current list of eight essential paraspeckle proteins includes: splicing factor proline- and glutamine-rich protein (SFPQ), the non-POU domain-containing octamer-binding protein (NONO), found in sarcoma (FUS), RNA binding protein 14 (RBM14), the Brahma-related gene-1 (BRG1), DAZ-associated protein 1 (DAZAP1), and two heterogeneous nuclear ribonucleoproteins, HNRNPK and HNRNPH3 (10, 12). Other proteins like PSP1 are enriched in PS but not required for their assembly (13).

Paraspeckles form co-transcriptionally. First, SFPQ and NONO load onto the nascent *Neat1*_2 transcript as it is being made, forming an intermediate *Neat1*_2 ribonucleoprotein. Then, FUS and RBM14 are recruited to drive the formation of mature paraspeckles (6, 10, 14). Paraspeckle-associated RBPs like SFPQ, NONO, FUS, HNRNPK, etc. have been implicated in processes like transcription, splicing, and polyadenylation SFPQ (15–19). As these and other proteins are incorporated into growing paraspeckles, their ability to participate in other nuclear gene expression pathways is altered. This sequestration mechanism has been proposed for SFPQ-mediated transcription of the RNA editing ADARB2 gene (20) and TDP-43 control of alternative polyadenylation (21). Likewise, association of the SWI/SNF complex protein ARID1B with the paraspeckle has been shown to influence alternative splicing in HEK293T cells (22).

Because of links between *Neat1* and cancer, neurodegenerative disease, and inflammatory disorders, there is growing interest in how *Neat1* and paraspeckles control cellular homeostasis and stress responses (2, 23). Specialized cell types, like neurons and immune cells, are constantly receiving and responding to environmental inputs that trigger remarkable changes to their transcriptomes and proteomes. Innate immune cells like macrophages are particularly exemplary of this behavior. Macrophages, our body’s first line of defense against foreign agents, express a panoply of pattern recognition receptors that allow them to sense pathogen- and damage-associated molecular patterns (PAMPs and DAMPs). When these sensors are engaged, they trigger a series of complex signal transduction cascades that activate transcription factors to turn on *de novo* expression of cytokines, chemokines, and antimicrobial mediators. Several studies have begun to link paraspeckles to innate immune gene expression and antiviral responses. We know that *Neat1*-deficient mice mount reduced inflammatory responses during models of peritonitis and pneumonia (24) and *Neat1* itself can be upregulated in response to DNA or RNA viral infection (25–27). Depletion of *Neat1* has been shown to promote expression of inflammatory genes like IL-8 via sequestration of repressive SFPQ from the IL-8 promoter (28) and has been linked to reduced dengue virus replication (29). Despite these intriguing links between paraspeckles and inflammation, it remains to be seen how paraspeckles function in *bona fide* immune cells like macrophages.

We hypothesized that nuclear MLOs like paraspeckles help the nucleus regulate transcription of innate immune genes in response to pathogen sensing. Consistent with a role for these MLOs in sensing and responding to pathogens, we found that paraspeckles undergo rapid changes in macrophages following several innate stimuli. Notably, we observed that paraspeckles aggregate almost immediately upon pattern recognition receptor engagement and disappear at 2h post-stimulation of TLR4, TLR2, or cGAS. We report that paraspeckle upregulation in macrophages requires active transcription and disassembly of paraspeckles/loss of *Neat1* RNA is dependent on the nuclear RNA exosome and the nuclear exosome targeting (NEXT) complex. In the absence of *Neat1*, both immortalized bone marrow derived macrophages (iBMDMs) and primary BMDMs fail to fully induce pro-inflammatory M1-associated genes and overexpress wound-healing M2-associated genes, resulting in uncontrolled bacterial and viral replication. These data uncover a critical role for *Neat1* and the paraspeckle in mounting a balanced innate immune gene expression program to control bacterial and viral replication in macrophages.

## RESULTS

### Paraspeckles are rapidly up- and down-regulated in response to innate agonist treatment of macrophages

To begin to define the dynamics of paraspeckle formation during macrophage activation, we employed a technique to simultaneously detect the *Neat1* lncRNA by RNA-FISH and paraspeckle proteins like PSP1 by immunofluorescence (FISH-IF). Our FISH probes only anneal to sequences in the paraspeckle-forming *Neat1_2* lncRNA, which we will refer to as *Neat1* from now on. Using FISH-IF for *Neat1* and PSP1, we observed that resting RAW 264.7 macrophages maintain two clear paraspeckles, consistent with previous reports demonstrating co-transcriptional paraspeckle formation at each *Neat1* genomic locus (30, 31). We then treated macrophages with lipopolysaccharide (LPS) (100 ng/ml) and performed FISH-IF over a time-course of treatment. LPS, a component of the outer membrane of gram-negative bacteria, stimulates expression of hundreds of innate immune genes (e.g., *Tnf* and *Il1b*) by engaging TLR4 and activating transcription factors like NFκB and IRF3 (**Fig. S1A**). At 0.5 hr post-LPS treatment, we observed a dramatic upregulation of paraspeckles in RAW 264.7 macrophages (**Fig. 1A**). At 1 hr, paraspeckle area remained high, but qualitatively, paraspeckles became more dispersed throughout the nucleus. Paraspeckle aggregation was concomitant with an approximately 2- to 3-fold increase in total *Neat1* RNA as measured by RT-qPCR (**Fig. 1B**; primers designed to amplify a region unique to the long *Neat1_2* isoform). At 2 hrs post-LPS, paraspeckle signal was virtually undetectable, with no observable *Neat1* or PSP1 puncta. Paraspeckle disaggregation was concomitant with a loss of *Neat1* RNA signal by RT-qPCR (**Fig. 1B**). Compared to our normal Zymo DirectZol column extractions, *Neat1_2* extraction efficiency improved with mechanical shearing or incubation at 55 deg (**Fig. S1B**), but the pattern of *Neat1_2* up- and down-regulation at each time point post-LPS treatment remained the same (**Fig. 1B** and **S1B**). By 4 hrs post-LPS, paraspeckles started to reform although total *Neat1* RNA remained low. At 6 hrs post-LPS, cells were heterogeneous in their paraspeckle numbers, with most cells having 0, 1, or 2 paraspeckles and about 20% of cells maintaining higher numbers (**Fig. S1C**). By 8h post-LPS, the percentage of cells maintaining >2 paraspeckles increased to ∼50%, suggestive of cells restarting the cycle of paraspeckle aggregation (**Fig. S1D**). Another abundant lncRNA, *Malat1*, which is encoded in the genome directly downstream of *Neat1,* showed no evidence of up or downregulation over an 8 hr LPS treatment (**Fig. S1E**). We observed similar paraspeckle dynamics in primary BMDMs (**Fig. 1C**), with paraspeckle area peaking at 0.5 hr post-LPS and signal ablation at 2 hrs post-LPS. Compared to reports in other cell types, maximum paraspeckle hyperaggregation following pathogen sensing in macrophages is incredibly fast (0.5 hr v. 3 hrs or more (28, 32)). To the best of our knowledge, the phenomena of paraspeckle disappearance at 2 hrs and subsequent recovery at 4 hrs post-LPS treatment has not been reported as part of normal cellular responses in other mammalian cell types, and thus might represent a macrophage-specific adaptation.

**Figure 1:**
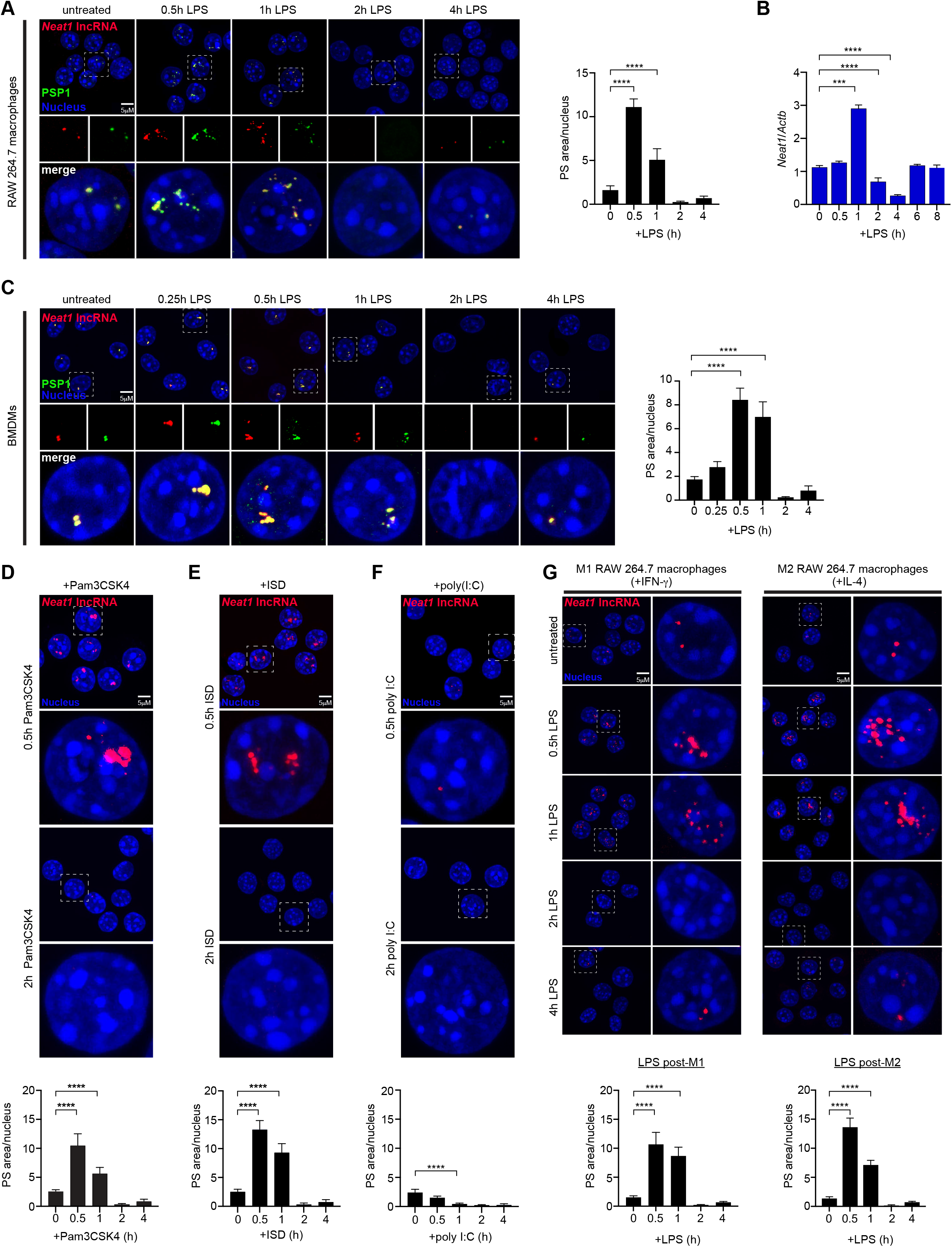
Nuclear paraspeckles are dynamically regulated following innate immune agonist treatment of macrophages. (A) RNA-FISH of *Neat1* (red) and immunofluorescence microscopy of PSP1 (green) in RAW 264.7 macrophages post-LPS treatment (100 ng/ml). Quantitation of paraspeckle area/nucleus on right. (B) RT-qPCR of *Neat1_2* transcript levels in RAW 264.7 macrophages post-LPS treatment (100ng/ml) shown relative to *Actb.* (C) As in (A) but with BMDMs treated with 10 ng/ml LPS. Quantitation of paraspeckle area/nucleus on right. (D) RNA-FISH of *Neat1* (red) in RAW 264.7 macrophages post-Pam3CSK4 treatment (100 ng/ml). Quantitation of paraspeckle area/nucleus below. (E) As in (D) but post-dsDNA transfection (ISD) (1 μg/ml). (F) As in (D) but post-dsRNA transfection (poly I:C) (500 ng/ml). (G) RNA-FISH of *Neat1* (red) in RAW 264.7 macrophages post-LPS treatment (100 ng/ml) following overnight polarization into M1- (+IFN-γ; 50 ng/ml) or M2 (+IL-4; 25 ng/ml)-like macrophages. Quantitation of paraspeckle area/nucleus below. <u>Statistical tests</u>: Data is presented as the mean of three biological replicates unless otherwise noted with error bars representing SEM. At least 100 cells were counted over multiple coverslips per condition. Statistical significance was determined using a one-way ANOVA (A-G). ***p<0.001, ****p<0.0001.

We next asked whether paraspeckle dynamics triggered by LPS, which activates pathogen sensing cascades via TLR4, were unique. Having seen very similar paraspeckle dynamics between RAW 264.7 macrophages cells and primary BMDMs, we chose to continue studies with the genetically tractable RAW 264.7 cell line. Likewise, having seen identical patterns for the *Neat1* lncRNA by FISH and the PSP1 protein by immunofluorescence microscopy across multiple experiments, we opted to track paraspeckles by *Neat1* FISH alone. Thus, we treated RAW 264.7 macrophages with a panel of innate immune agonists and measured paraspeckle formation by *Neat1* RNA-FISH. Treatment of cells with the TLR2 agonist Pam3CSK4 (**Fig. 1D**; **Fig. S1F**) or the cGAS agonist cytosolic dsDNA (ISD) (**Fig. 1E**; **Fig. S1G**) triggered paraspeckle dynamics that closely followed those induced by LPS. Surprisingly, transfection of poly I:C, a dsRNA agonist of TLR3 (endosomal) and RIG-I/MDA5 (cytosolic) RNA sensing cascades, rapidly ablated *Neat1* signal in the nucleus (**Fig. 1F**; **Fig. S1H**). Our poly I:C results contrast a previous report that showed paraspeckle upregulation in HeLa cells post-poly I:C transfection (33), further arguing that that macrophages regulate paraspeckles in a fundamentally different way than non-immune cells.

Macrophage polarization is an important determinant in dictating innate immune outcomes (34). Macrophages can take on a classical pro-inflammatory M1 state when treated with IFN-γ or can be alternatively activated to a wound-healing M2 state after treatment with IL-4. To determine whether macrophage polarization impacts paraspeckle dynamics, we treated RAW 264.7 macrophages overnight with IFN-γ (M1) or IL-4 (M2) and confirmed polarization by measuring canonical M1/M2 transcripts by RT-qPCR (**Fig. S1I**). We did not see dramatic upregulation of PS at 0.5 hr post-IFN-γ or IL-4 treatment alone, suggesting that treatment with these cytokines is not sufficient to upregulate paraspeckles (**Fig. S1J**). We then repeated our LPS time-course in these M1 or M2 macrophages. In both cases, we qualitatively observed hyper-accumulation of *Neat1* by RNA-FISH at 0.5 and 1 hr, suggesting that polarized macrophages are primed to upregulate paraspeckles (**Fig. 1G**). Downregulation of paraspeckles occurred with similar kinetics in M1 and M2-polarized macrophages (**Fig. 1G**). Together, these data identify nuclear paraspeckles as immune-responsive MLOs in macrophages that are dynamically regulated downstream of multiple innate sensing cascades.

### Paraspeckle upregulation in macrophages can sequester nuclear RNA binding proteins

Paraspeckles contain many copies of the *Neat1* RNA and an array of RNA binding proteins (35). To better understand the nature of the macrophage paraspeckle, we next asked whether canonical paraspeckle proteins localize with *Neat1* in the macrophage nucleus and whether these associations are altered by LPS treatment. Consistent with reports in other cell types, we observed significant colocalization between *Neat1* and PSP1 (**Fig. 2A**), SFPQ (**Fig. 2B**), and FUS (**Fig. 2C**) in both resting and LPS-stimulated macrophages. Interestingly, non-*Neat1*-colocalizing FUS puncta were also upregulated in response to LPS, suggesting that macrophage activation triggers FUS sequestration into non-paraspeckle compartments as well. Association between NONO and *Neat1* appeared to follow distinct dynamics, with approximately 30% more colocalization measured in LPS-treated compared to resting macrophages (**Fig. 2D**). As total cellular levels of these paraspeckle proteins remained constant over a 1 hr time-course of LPS treatment, these data support a model whereby already synthesized paraspeckle proteins in the nucleus are brought into paraspeckles as they aggregate (**Fig. S2A**). Components of the SWI/SNF nucleosome remodeling complex, which play an important role in activating expression of secondary response genes like *Il6* (36) have been found in paraspeckles in other model cell types. Surprisingly, BRG1 and BRM, deemed essential for paraspeckle assembly in HeLa cells (37), did not display obvious punctate staining reminiscent of paraspeckles pre- or post-LPS treatment (**Fig. 2E and S2B**), despite clear aggregation of BRM in resting macrophages. Lack of evidence for BRM and BRG1 enrichment in macrophage paraspeckles begins to suggest that the composition of these condensates may be cell-type specific.

**Figure 2:**
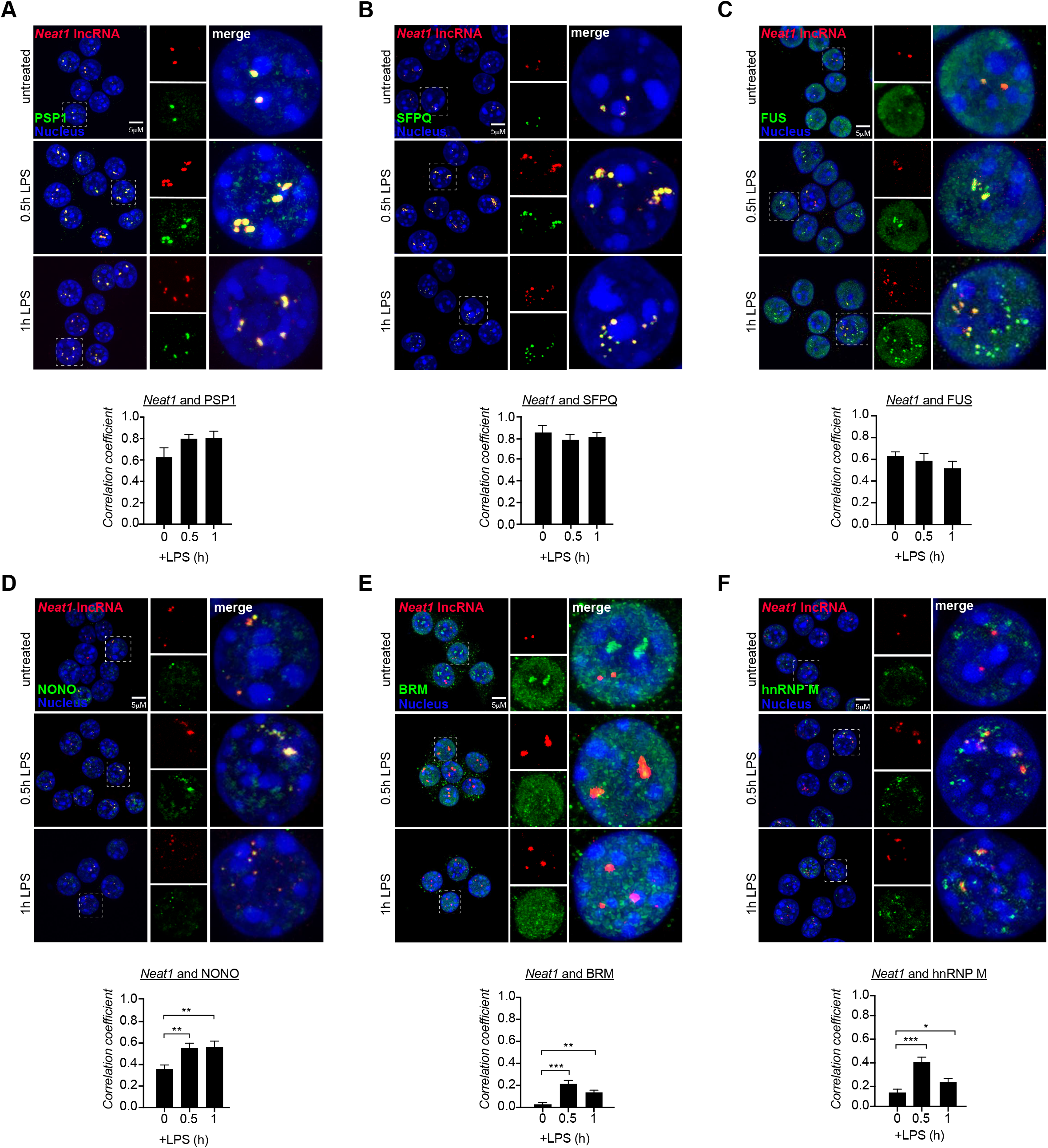
Paraspeckle upregulation in macrophages sequesters nuclear RNA binding proteins. (A) RNA-FISH of *Neat1* (red) and immunofluorescence microscopy of PSP1 (green) in RAW 264.7 macrophages at 0.5 hr and 1 hr post-LPS treatment (100 ng/ml). Correlation coefficient between *Neat1* and PSP1 quantified below. (B) As in (A) but for SFPQ. (C) As in (A) but for FUS. (D) As in (A) but for NONO. (E) As in (A) but for BRM. (F) As in (A) but for hnRNP M. <u>Statistical tests</u>: Data is presented as the mean of three biological replicates unless otherwise noted with error bars representing SEM. At least 100 cells were counted over multiple coverslips per condition. Colocalization coefficient was measured using the ImageJ plugin Coloc2. Statistical significance was determined using a one-way ANOVA (A-F). *p<0.05, **p<0.01, ***p<0.001.

As over 20 RBPs involved in a variety of nuclear processes (pre-mRNA splicing, RNA editing, mRNA export, etc.) have been shown to localize to and/or purify with the paraspeckle in non-immune cells (32), we posited that RBPs and immune-associated proteins could be brought into paraspeckles during macrophage activation. One RBP that reportedly interacts with paraspeckle proteins and has links to innate immune gene expression is the splicing factor hnRNP M, which represses intron removal in inflammatory cytokine pre-mRNAs like *Il6* (38). By FISH-IF, we saw a marked increase in colocalization between hnRNP M and *Neat1* at 0.5 hr post-LPS treatment (**Fig. 2F**). Because it would be an obvious mechanism through which paraspeckles could influence macrophage gene expression, we also tested whether paraspeckles sequester innate immune transcription factors that are activated downstream of pattern recognition receptors like TLR4. Overall, colocalization between *Neat1* and the two factors queried, NFκB and STAT1, was low (**Fig. S2C-D**), although we did measure a slight, but statistically significant, increase in colocalization between *Neat1* and NFκB in LPS-treated macrophages. This could represent enrichment of NFκB at the site of *Neat1* transcription and would be consistent with ChIP-seq experiments that show enrichment of the NFκB subunit RelA at the *Neat1* promoter in response to LPS (**Fig. S3D**). Together, these data hint at compositional differences between macrophage paraspeckles and those previously described in other model cell types. They also suggest that RBPs previously linked to post-transcriptional regulation of innate immune gene expression (e.g., hnRNP M) have links to the paraspeckle in macrophages.

### Transcription and MAPK signaling are required to maintain and upregulate paraspeckles in macrophages

Given the rapid up- and downregulation of paraspeckles during the early macrophage response to LPS, we set out to investigate the cellular pathways that control paraspeckle maintenance and aggregation in these cells. First, we asked whether transcription was required for paraspeckle upregulation after LPS treatment. Briefly, RAW 264.7 macrophages were treated with the transcription inhibitor actinomycin D (ActD) for 0.5 hr and *Neat1* was monitored by FISH and RT-qPCR 0.5 hr post-LPS. Treatment with ActD not only prevented LPS-induced paraspeckle upregulation but also inhibited paraspeckle maintenance all together, as we could no longer detect *Neat1* puncta in resting macrophages after 0.5 hr of ActD (**Fig. 3A**). Curiously, this loss of paraspeckle signal did not correspond directly with total cellular levels of *Neat1* lncRNA, which remained high for at least 1 hr after ActD (**Fig. 3B**) and ActD/LPS treatment (**Fig. 3C**). This suggests that while active transcription is needed to maintain and aggregate paraspeckles, this is likely due more to reliance on the act of transcription for assembling paraspeckles, rather than a need to synthesize new *Neat1*. The kinetics of *Neat1* lncRNA turnover after transcription shut off were on the order of 2 hrs in RAW 264.7 macrophages (**Fig. 3B-C; Fig. S3A**)—the same time-point at which we see loss of paraspeckles after LPS treatment (**Fig. 1A**). Degradation of the *Neat1* RNA after transcriptional shut-off far exceeded that of a stable housekeeping gene like *Gapdh* (**Fig. 3D**; **Fig. S3B**) and was even faster than that of another previously identified short lived non-coding RNA, *Kcnq10t1* (39) (**Fig. 3E**; **Fig. S3C**), consistent with earlier reports of *Neat1* being an unstable/short-lived RNA (39). Together, these data argue that in macrophages *de novo* transcription is required both to maintain paraspeckles, as was previously reported in myoblasts (31), and to aggregate paraspeckles upon LPS treatment.

**Figure 3:**
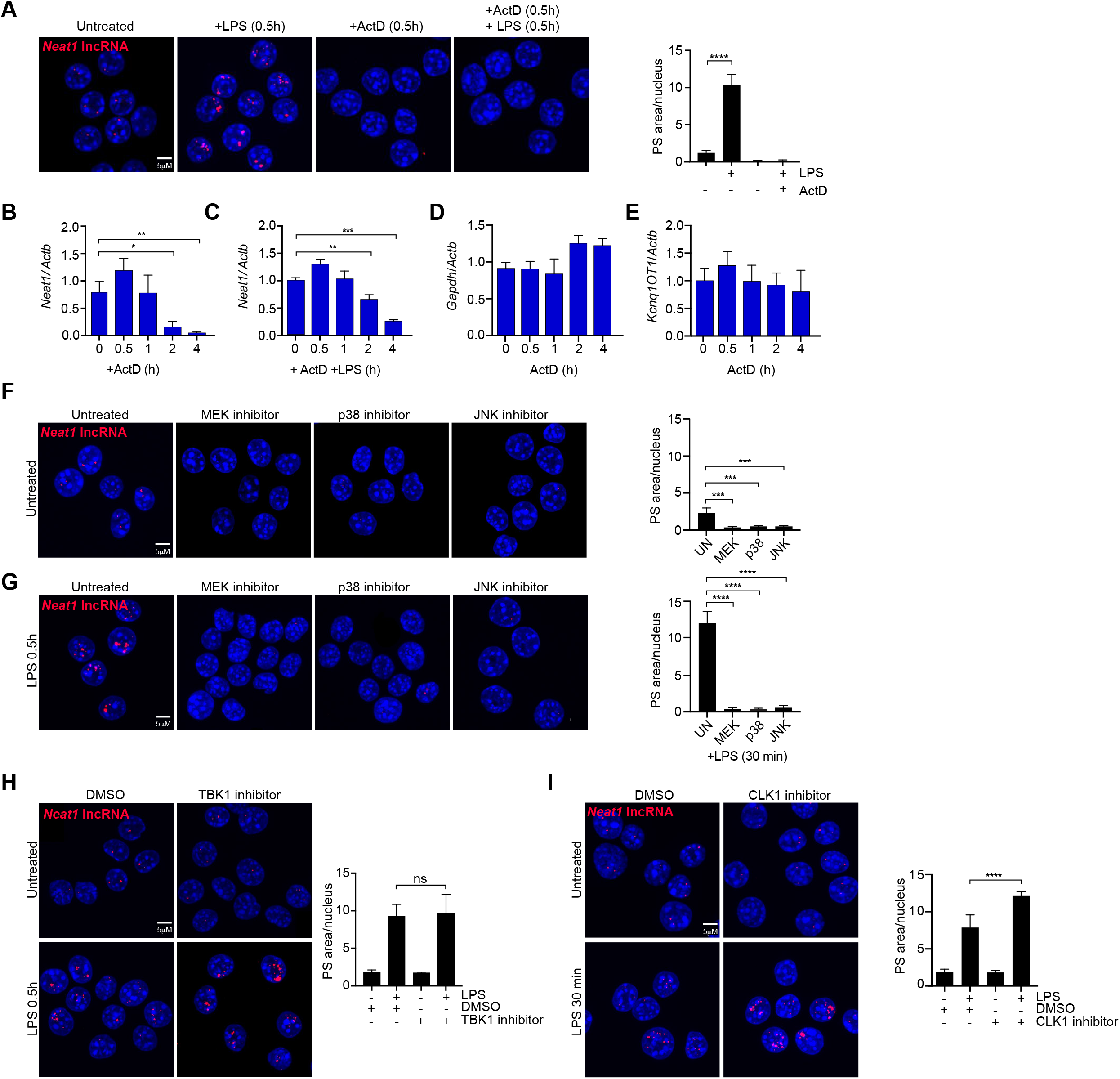
*De novo* transcription and basal MAPK signaling maintain paraspeckles in macrophages. (A) RNA-FISH of *Neat1* (red) in RAW 264.7 macrophages after actinomycin D treatment (ActD) (5μg/ml) followed by LPS stimulation for timepoints indicated. Quantitation of paraspeckle area/nucleus on right. (B) RT-qPCR of *Neat1_2* transcript levels in RAW 264.7 macrophages after ActD treatment, (5μg/ml) shown relative to *Actb*. (C) As in (B) but with LPS (100 ng/ml) treatment following 0.5 hr ActD. (D) As in (B) but for *Gapdh*. (E) As in (B) but for *Kcnq1OT1*. (F) RNA-FISH of *Neat1* (red) in RAW 264.7 macrophages post-MAPK inhibitor treatment (MEK inhibitor (U0126; 25 μM), JNK inhibitor (SP600125; 25 μM), p38 inhibitor (SB203580; 10 μM)) for 0.75 hr. Quantitation of paraspeckle area/nucleus on right. (G) As in (F) but with the addition of LPS (100 ng/ml) for 0.5 hr following ActD. Quantitation of paraspeckle area/nucleus on right. (H) RNA-FISH of *Neat1* (red) in RAW 264.7 macrophages after treatment with the TBK1 inhibitor (GSK-8612; 10 μM) at 0 and 0.5 hr post-LPS stimulation. Quantitation of paraspeckle area/nucleus on right. (I) As in (H) but with CLK1 inhibitor (Cpd 23; 10 μM). Quantitation of paraspeckle area/nucleus on right. <u>Statistical tests</u>: Data is presented as the mean of three biological replicates unless otherwise noted with error bars representing SEM. At least 100 cells were counted over multiple coverslips per condition. Statistical significance was determined using two-tailed Student’s t-test (B-E) or one-way ANOVA (A, F-I). *p<0.05, **p<0.01, ***p<0.001, ****p<0.0001.

We next sought to identify the signaling cascades that promote *Neat1* transcription and upregulation upon LPS treatment. LPS stimulation of TLR4, and TLR signaling in general, activate a complex network of kinase cascades, including MEK, JNK, and p38 MAP kinases, AKT and PI3-K, IκB kinase, and the noncanonical IκB kinase homologs IKK-ε and TBK1 (40, 41). To begin to test the role of MAPKs in regulating PS dynamics in macrophages, we LPS-treated RAW 264.7 macrophages for 0.5 hr following pretreatment for 0.75 hr with a MEK inhibitor (U0126), a JNK inhibitor (SP600125), or a p38 inhibitor (SB203580). Remarkably, not only did we see paraspeckles fail to accumulate in response to LPS after all three MAPK inhibitor treatments, but we also saw loss of paraspeckles in untreated macrophages, suggesting that basal MAPK signaling is important for maintaining paraspeckles in resting cells (**Fig. 3F-G**). Inhibiting other potentially relevant cellular kinases like TBK1 (**Fig. 3H, S3F**), which is activated down-stream of TLR4 to phosphorylate the transcription factor IRF3, or CLK1, which is activated downstream of AKT signaling to regulate phosphorylation of SR proteins (42, 43), did not ablate paraspeckles (**Fig. 3I-S3G**). In fact, treatment with the CLK1 inhibitor resulted in modest hyperaggregation of *Neat1* at 0.5 hr post-LPS treatment. These results implicate MAPKs as positive regulators of paraspeckle maintenance and hint at SR protein phosphorylation negatively regulating paraspeckle aggregation in macrophages.

### The *Neat1* lncRNA is targeted to the nuclear exosome by the NEXT complex to regulate PS dynamics in macrophages

In the nucleus, RNA turnover is controlled by the RNA exosome, a multiprotein complex responsible for 3’ end processing and/or degradation of RNAs (44). The exosome forms a barrel structure and has two associated 3’ to 5’ exoribonucleases: EXOSC10/RRP6 and DIS3 (**Fig. 4A**). RNAs are targeted to the exosome for processing or degradation by one of three accessory protein complexes: NEXT, TRAMP, or PAXT/PCC. NEXT is involved in turnover of introns released by pre-mRNA splicing and unstable RNAs from pervasive transcription. TRAMP degrades RNAs like pre-rRNAs, cryptic unstable transcripts, as well as a variety of aberrant small RNAs (tRNAs, ncRNAs, snoRNAs, snRNAs). PAXT/PCC is responsible for bringing nuclear ncRNAs with long polyA tails to the exosome. To begin to implicate the exosome in regulation of *Neat1* and paraspeckles in macrophages, we transfected siRNAs designed against *Mtr4,* alongside a non-targeted control (NC), into RAW 264.7 macrophages. MTR4 is a member of the SKI2 family of RNA helicases that is common to all three nuclear exosome targeting complexes. At 48 hours post-transfection, we achieved >90% knockdown of *Mtr4* (**Fig. S4A**). By RNA-FISH, we observed a dramatic increase in *Neat1* signal in resting *Mtr4* knockdown macrophages compared with NC siRNA control cells (**Fig. 4B**). This correlated with total *Neat1_2* transcript accumulation (**Fig. 4C**) and is consistent with previous reports of *Neat1* instability and a role for the exosome in controlling *Neat1* turnover (45). *Mtr4* knockdown did not result in accumulation of other abundant nuclear mRNAs or snRNAs (**Fig. S4B**).

**Figure 4:**
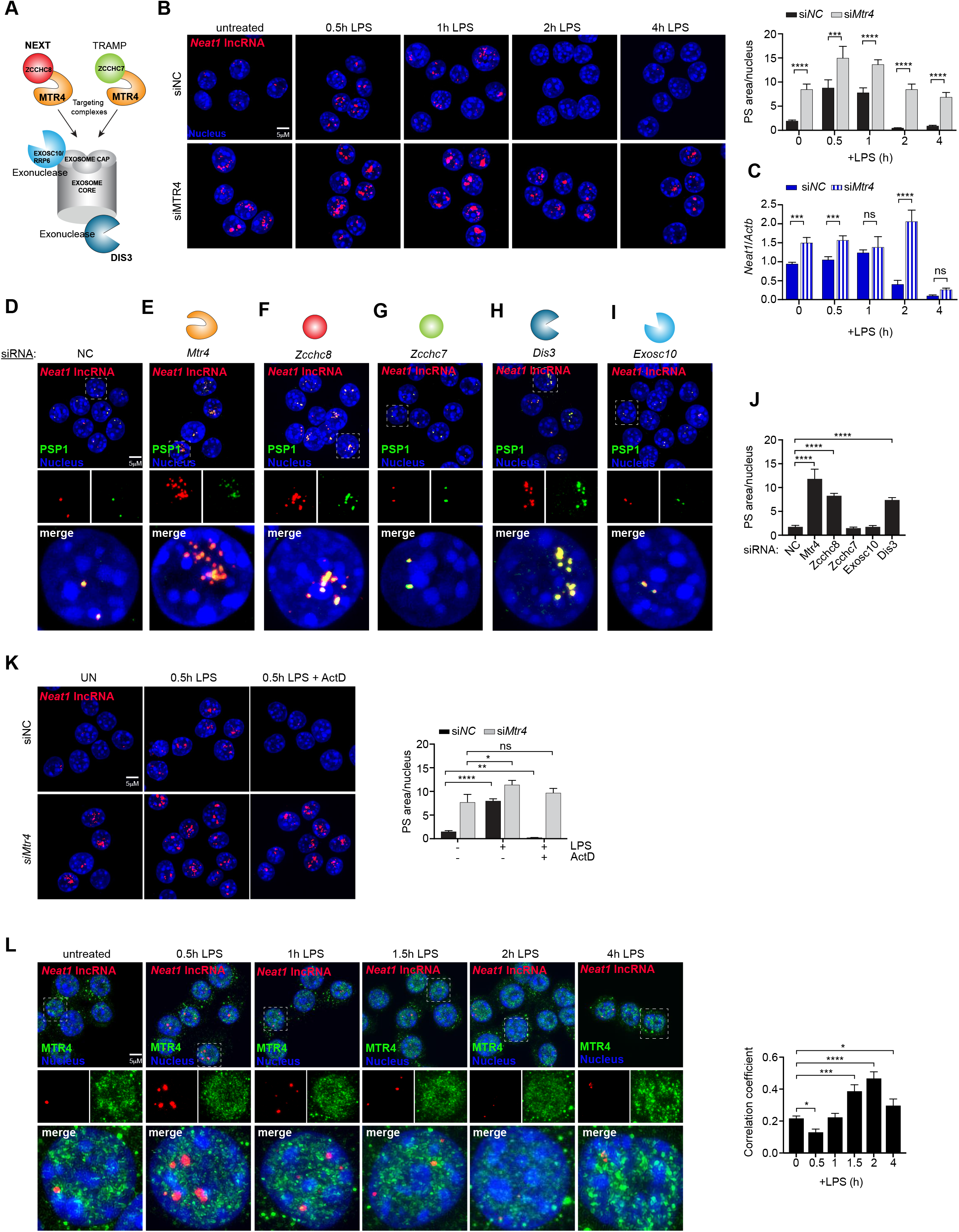
The NEXT complex targets the *Neat1* lncRNA to the nuclear exosome to regulate paraspeckle dynamics in macrophages. (A) Model of nuclear RNA exosome and the NEXT and TRAMP targeting complexes. Colored shapes denote factors knocked down in panels E-I. (B) RNA-FISH of *Neat1* (red) in *Mtr4* knockdown RAW 264.7 macrophages 48 hrs after Silencer Select siRNA transfection, alongside a negative control (siNC), over a time-course of LPS treatment (100 ng/ml). Quantitation of paraspeckle area/nucleus on right. (C) RT-qPCR of *Neat1_2* transcript levels in *siMtr4* and *siNC* RAW 264.7 macrophages post-LPS treatment (100ng/ml) shown relative to *Actb.* (D) RNA-FISH of *Neat1* (red) and immunofluorescence microscopy of PSP1 (green) in *siNC* RAW264.7 macrophages. (E) As in (D), but in *siMtr4*-treated RAW 264.7 macrophages. (F) As in (D), but in *siZcchc8*-treated macrophages. (G) As in (D), but in *siZcchc7*-treated macrophages. (H) As in (D), but in *siDis3*-treated macrophages (I) As in (D), but in *siExosc10*-treated macrophages. (J) Quantitation of paraspeckle area/nucleus of 3D-I. (K) RNA-FISH of *Neat1* (red) in *siNC* and *siMtr4*-treated RAW 264.7 macrophages in untreated cells, +LPS (100 ng/ml; 0.5 hr) or +ActD (5 mg/ml; 0.5 hr) followed by LPS (100 ng/ml; 0.5 hr). (L) RNA-FISH of *Neat1* (red) and immunofluorescence microscopy of MTR4 (green) in RAW 264.7 macrophages post-LPS treatment (100 ng/ml). Quantitation of paraspeckle area/nucleus on right. <u>Statistical</u> <u>tests</u>: Data is presented as the mean of three biological replicates unless otherwise noted with error bars representing SEM. At least 100 cells were counted over multiple coverslips per condition. Colocalization coefficient was measured using the ImageJ plugin Coloc2. Statistical significance was determined using a Student’s t-test (A-C) or one-way ANOVA (J-L). *p<0.05, **p<0.01, ***p<0.001, ****p<0.0001.

*Neat1* aggregates that form in the absence of MTR4 are *bona fide* structured PSs, as they each show significant enrichment with the paraspeckle protein PSP1 (**Fig. 4D-E**). Even though *Mtr4* KD macrophages have high numbers of paraspeckles at rest, these numbers still increase after LPS treatment (*siMtr4* time 0 vs. 0.5 hr vs. 1h post-LPS; **Fig. 4B**). We interpret this to mean that aggregation of paraspeckles following LPS stimulation occurs independently of the exosome. We can, however, implicate the exosome in paraspeckle disassembly at 2 hrs post-LPS, as we do not see the characteristic loss of *Neat1* signal at 2 hrs post-LPS in *Mtr4* KD macrophages (**Fig 4B**). While bulk measurements of *Neat1_2* cellular transcripts reflect this phenotype at 2 hrs (i.e. *Neat1* is higher in *siMtr4* relative to siNC cells), by 4 hrs, *Neat1_2* levels are very low regardless of whether cells have MTR4. This discrepancy at 4 hrs may stem from incomplete knockdown of *Mtr4,* where our bulk measurements reflect a mixture of cells, some of which have *Mtr4* knocked down and others that have normal levels of *Mtr4*. It is also possible that some kind of transcriptional shut-off limits *Neat1* abundance even in exosome-knockdown cells. Regardless, we can conclude from these data that TLR4 engagement signals exosome-mediated dissolution of paraspeckles and turnover of *Neat1* at 2 hrs post-LPS treatment in macrophages.

Having implicated the MTR4 RNA exosome helicase in *Neat1* turnover and paraspeckle aggregation (**Fig. 4B-E**), we next sought to pin down the nature of the targeting complex and exonuclease that regulate *Neat1* stability in macrophages. Since *Neat1_2* is not polyadenylated, we ruled out a role for the PAXT/PCC targeting complex. To implicate either NEXT or TRAMP in *Neat1* turnover, we transfected RAW 264.7 macrophages with siRNAs directed against *Zcchc8* (NEXT) (**Fig. S4C**) and *Zcchc7* (TRAMP) (**Fig. S4D**) and observed paraspeckles by IF-FISH in resting cells. Loss of *Zcchc8* upregulated paraspeckles in a similar fashion to loss of *Mtr4* (**Fig. 4E, F, J**), while loss of *Zcchc7* had no impact on paraspeckle number or size (**Fig. 4G, J**). From this, we can conclude that *Neat1* is likely targeted to the exosome in macrophages by the NEXT complex. We performed a similar experiment to determine the exoribonuclease that degrades *Neat1*, comparing paraspeckles in resting *Exosc10* versus *Dis3* knockdown macrophages (**Fig. S4E-F**). We observed a clear upregulation of paraspeckles in cells transfected with siRNAs directed against *Dis3* but not *Exosc10* (**Fig. 4H-J**). Together, these data demonstrate that in resting macrophages, *Neat1* stability and paraspeckle maintenance is controlled by the NEXT targeting complex and the DIS3 exoribonuclease.

Next, we asked whether co-transcriptional PS assembly and exosome targeting of *Neat1* in macrophages are separable mechanisms. To do so, we treated *siNC-* and *siMtr4-transfected* macrophages with LPS in the presence or absence of ActD, as in Fig. 3A. Whereas ActD completely ablated *Neat1* signal after 0.5 hr in siNC cells (as we observed for wild-type macrophages in Fig. 3A), ActD had no impact on paraspeckles in *siMtr4* KD cells (**Fig. 4K**). Therefore, blocking the exosome allows for paraspeckle maintenance even in the absence of *de novo* transcription, suggesting a tug-of-war between transcription and exosome turnover in maintaining *Neat1* and paraspeckles in macrophages.

Lastly, we supposed that if *Neat1* is constitutively degraded by the exosome, as is suggested by the high number of paraspeckles we see in resting *Mtr4* knockdown macrophages (**Fig. 4B**), we might be able to detect co-localization between exosome components and the *Neat1* lncRNA. To test this, we performed FISH-IF using antibodies against MTR4. Indeed, we can detect colocalization between *Neat1* aggregates and MTR4 in resting cells, consistent with constitutive turnover of *Neat1* by the exosome. Moreover, over the course of LPS stimulation, we saw an inverse relationship between MTR4-*Neat1* colocalization and paraspeckle aggregation, whereby colocalization was lowest at 0.5 hr, when paraspeckles are growing, and highest at 1-2 hrs when paraspeckles disintegrate (**Fig. 4L**). These data suggest a regulated mechanism of paraspeckle targeting to the exosome in macrophages and hint at undescribed links between pattern recognition receptor engagement and exosome activity.

### *Neat1* is required to activate the innate immune response in macrophages

Having observed dramatic, regulated reorganization of paraspeckles in macrophages following LPS treatment, we finally asked whether ablating *Neat1*, and disrupting this paraspeckle cycle, impacts the macrophage innate immune response. To this end, we acquired mice that do not express *Neat1* due to incorporation of a lacZ cassette at the 5’ end of the *Neat1* gene (46) (**Fig. S5A)**. From these *Neat1* KO mice, we differentiated BMDMs, treated them with LPS (10ng/ml) and isolated RNA for high-throughput sequencing and differential expression analysis at 0, 2 hrs and 4 hrs post-stimulation, to capture the early transcriptional response to LPS. Loss of *Neat1* in these cells was confirmed by RNA-FISH (**Fig. 5A**). To enable identification of non-coding RNAs and incompletely processed RNAs whose abundance may be altered in the absence of *Neat1*, we generated our sequencing libraries using ribodepletion. Differential expression analysis uncovered hundreds of misregulated genes in *Neat1* KO macrophages (log_2_FC > 0.5, < -0.5; adj. p-value <0.05) (208 genes at rest, 348 genes at 2 hrs post-LPS, and 352 genes at 4 hrs post-LPS) (**Supplemental Table 1** and **Fig. S5B**). Approximately one-third of *Neat1*-dependent genes reach this differential expression threshold in all three conditions (i.e. 0, 2, 4 hrs post-LPS) (**Fig. S5B**). Loss of *Neat1* resulted in both up and downregulation of gene expression with up- versus down-regulated genes segregating into distinct cellular pathways. Specifically, Ingenuity Pathway Analysis identified upregulation of genes with functional connections to hepatic fibrosis, pulmonary fibrosis, and wound healing in *Neat1* KO macrophages (**Fig. 5B**). Curiously, these are functions typically attributed to alternatively activated or M2-like macrophages that specialize in cellular proliferation and tissue repair (**Fig. 5D**). Many genes in these categories are normally turned off as part of the pro-inflammatory macrophage response to LPS (**Fig. S3C**). Genes that were downregulated in *Neat1* KO macrophages are enriched in pathways related to interferon signaling, hypercytokinemia in influenza pathogenesis, pathogen induced cytokine storm signaling, and pattern recognition receptor activation (**Fig. 5C, E-F**). These pathways are related to pro-inflammatory or M1-like responses in macrophages and are comprised of genes that are turned on in response to LPS. As we observe *Neat1*-dependent differences in reads aligning to both intronic and exonic sequences, our results suggest a defect in transcription and not pre-mRNA splicing (**Fig. 5G-H**).

**Figure 5:**
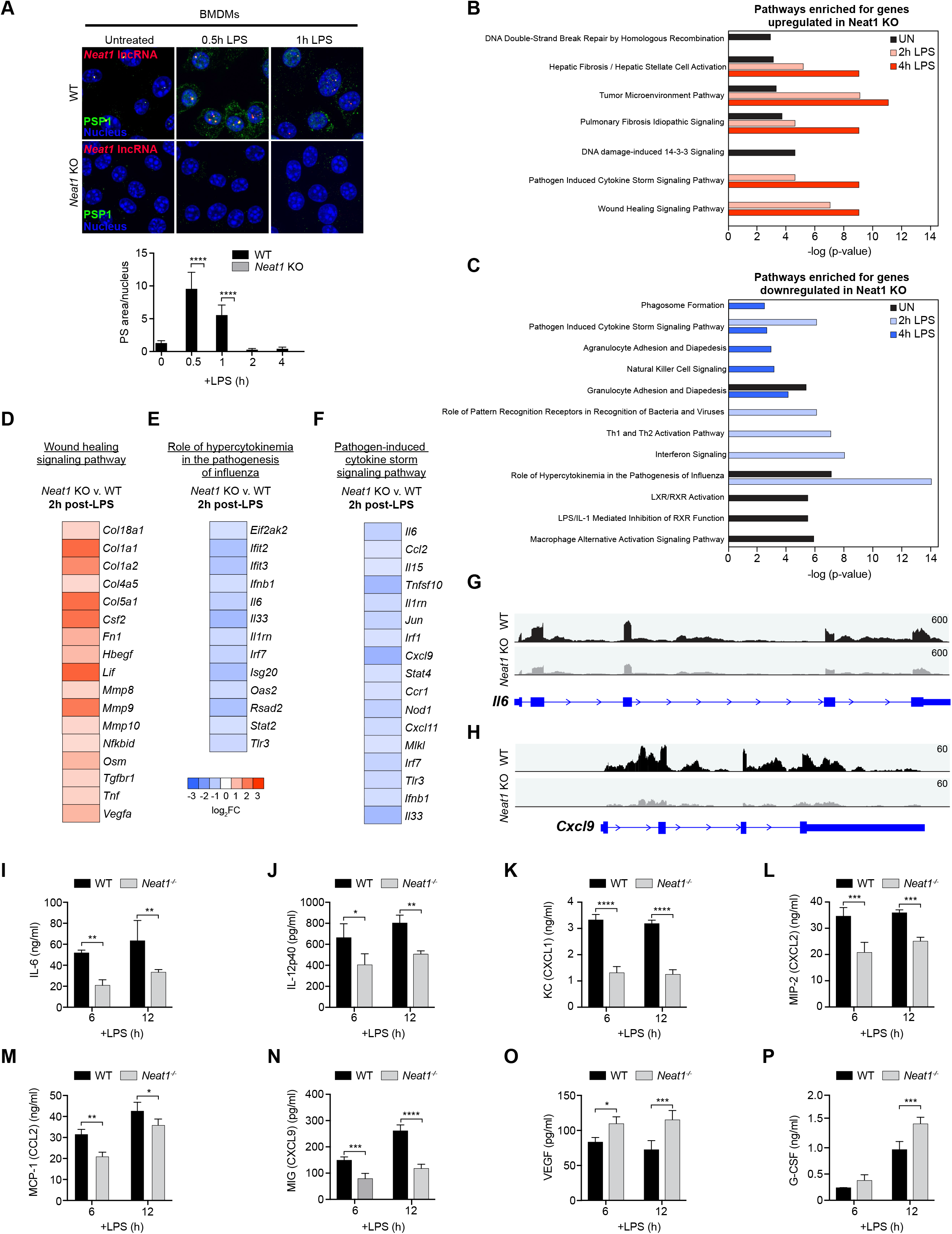
*Neat1* is required for proper up and downregulation of innate genes during the early macrophage response to LPS. (A) RNA-FISH of *Neat1* (red) and immunofluorescence microscopy of PSP1 (green) in WT and *Neat1* KO BMDMs over a 1 hr time-course of LPS treatment (10ng/ml). Quantitation of paraspeckle area/nucleus over full 4 hr time course below. (B) Ingenuity pathway analysis (IPA) for differentially expressed upregulated genes in *Neat1* KO vs. WT BMDMs (log_2_FC > 0.5; p<0.05) at 0, 2, and 4 hrs post-LPS treatment (10 ng/ml). (C) As in (B) but for downregulated genes. (D) Log_2_FC of differentially expressed genes in *Neat1* KO BMDMs categorized as “wound healing signaling pathway” by IPA. (E) As in (D) but for “role of hypercytokinemia in the pathogenesis of influenza.” (F) As in (E) but for “pathogen-induced cytokine storm signaling pathway.” (G) Integrated Genomics Viewer tracks of RNA-seq reads from a representative WT and *Neat1* KO sample at the *Il6* genomic locus (mm10). (H) As in (G) but for *Cxcl9*. (I) Measurements of IL-6 levels in the supernatants of WT and *Neat1* KO BMDM at 6 and 12 hrs post-LPS treatment (10 ng/ml) via cytokine array. (J) As in (I) but for IL-12p40. (K) As in (I) but for KC (CXCL1). (L) As in (I) but for MIP-2 (CXCL2). (M) As in (I) but for MCP-1 (CCL2). (N) As in (I) but for MIG (CXCL9). (O) As in (I) but for VEGF. (P) As in (I) but for G-CSF. <u>Statistical tests</u>: Data is presented as the mean of four biological replicates unless otherwise noted with error bars representing SEM. Statistical significance in (A and I-P) was determined using a Student’s t-test. *p<0.05, **p<0.01, ***p<0.001, ****p<0.0001.

To determine whether defective innate immune transcript accumulation in *Neat1* KO macrophages is borne out at the level of cytokine and chemokine secretion, we collected supernatants from WT and *Neat1* KO BMDMs and measured cytokine and chemokine secretion by multiplex cytokine array (Eve Technologies). Although we saw little evidence for differences in secretion of cytokines in resting *Neat1* KO and WT BMDMs (**Fig. S5D-M**), we observed defective expression/secretion of several cytokines and chemokines at 6 and 12 hrs post-LPS treatment in *Neat1* KO BMDMs. Factors that were downregulated by loss of *Neat1*, such as IL-6, KC/CXCL1, MIP-2/CXCL2, and MCP-1/CCL2, play critical roles in promoting inflammation and infiltration of neutrophils and lymphocytes to sites of infection (**Fig. 5I-N**). Cytokines upregulated by loss of *Neat1*, such as VEGF (encoded by the *Vegfa* gene) and G-CSF (encoded by the *Csf3* gene), are involved in cell proliferation and differentiation (**Fig. 5O-P**). Taken together, these data implicate *Neat1* in two critical aspects of the macrophage response to LPS: eliciting a balanced inflammatory response and differentiation into classical vs. alternative activation states.

### *Neat1* CRISPRi iBMDMs fail to execute antibacterial and antiviral defenses

As a strategy to overcome breeding limitations of the *Neat1* KO mice (47) and to expand upon our findings in primary cells, we designed a CRISPR interference-based approach to ablate *Neat1* from immortalized bone marrow derived macrophages. Using lentiviral transduction, we introduced a guide RNA directed against the *Neat1* promoter, alongside an untargeted control, into immortalized BMDMs expressing an endonuclease-deficient form of Cas9 (deactivated (dCas9) (48). We confirmed loss of *Neat1* expression in these CRISPRi cell lines by RT-qPCR (**Fig. S6A**) and by *Neat1* FISH (**Fig. 6A**). Over a 4 hr LPS treatment, paraspeckle dynamics in iBMDMs generally mirrored those that we report in RAW 264.7 macrophages and BMDMs, although we generally observed more cell-to-cell variation in paraspeckle number in the off-target gRNA-expressing iBMDMs.

**Figure 6:**
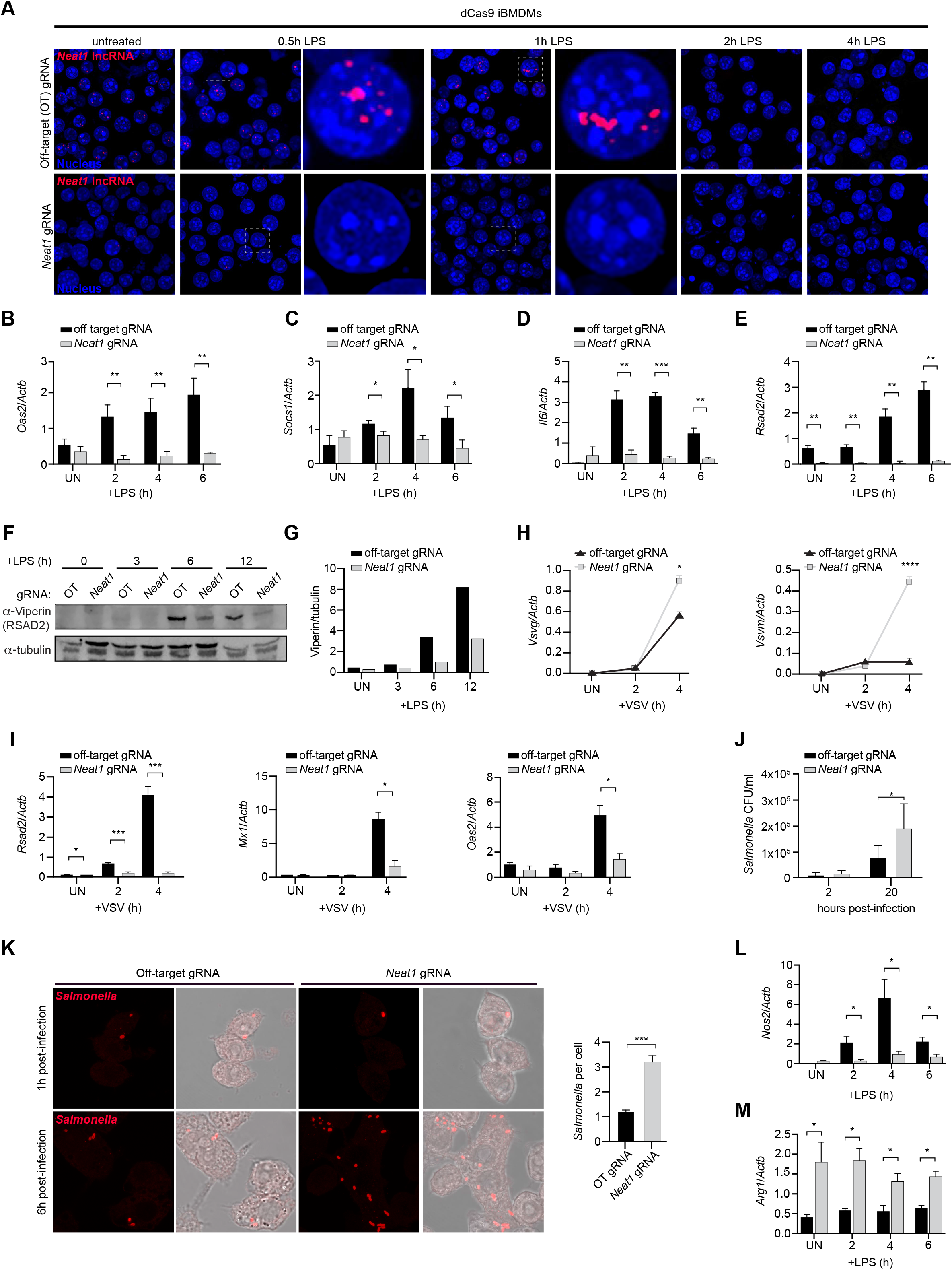
*Neat1* is required for antimicrobial responses in macrophages. (A) RNA-FISH of *Neat1* (red) at 0, 0.5, 1, 2, and 4 hrs post-LPS treatment (100ng/ml) of dCas9 iBMDMs transduced with an off-target (OT) gRNA lentiviral construct or a *Neat1* promoter-targeted gRNA lentiviral construct. Zoom in shown for 0.5 and 1 hr time points. (B) RT-qPCR of *Oas2* transcript levels in off-target gRNA and *Neat1* gRNA dCas9-expressing iBMDMs post-LPS treatment (100 ng/ml), shown relative to *Actb.* (C) As in (B), but for *Socs1*. (D) As in (B), but for *Il6*. (E) As in (B), but for *Rsad2*. (F) Immunoblot of Viperin (encoded by *Rsad2*) in off-target gRNA and *Neat1* gRNA dCas9-expressing iBMDMs post-LPS treatment (100 ng/ml). Quantitation, relative to tubulin, shown on right. (G) RT-qPCR of *Vsvg* and *Vsvm* transcript levels in off-target gRNA and *Neat1* gRNA dCas9-expressing iBMDMs after VSV infection (MOI = 1) for 0, 2 and 4 hrs, shown relative to *Actb*. (H) As in (G) but measuring macrophage *Rsad2*, *Mx1*, and *Oas2* transcript levels. (I) Colony-forming units (CFUs) of *Salmonella* recovered from off-target gRNA and *Neat1* gRNA dCas9-expressing iBMDMs at 2 and 20 hrs post-infection (MOI = 10). (J) Immunofluorescence of *Salmonella* (anti-LPS antibody; red) at 1 and 6 hrs post-*Salmonella* infection (MOI=10) in off-target gRNA and *Neat1* gRNA dCas9-expressing iBMDMs. (K) Quantification of *Salmonella* bacilli in (J). (L) RT-qPCR of *Nos2* transcript levels in off-target gRNA and *Neat1* gRNA dCas9-expressing iBMDMs at 2, 4, and 6 hrs post-LPS (100 ng/ml), shown relative to *Actb*. (M) As in (L) but for *Arg1*. <u>Statistical tests</u>: Data is presented as the mean of three biological replicates unless otherwise noted with error bars representing SEM. In J, over 100 cells were counted over multiple coverslips for each condition. Statistical significance was determined using a Students t-test (B-G, J, L-M). *p<0.05, **p<0.01, ***p<0.001.

Consistent with our RNA-seq data collected from BMDMs, many LPS-induced genes, including *Oas2*, *Socs1*, *Il6*, and *Rsad2*, failed to be effectively upregulated in *Neat1* gRNA iBMDMs (**Fig. 6B-E**). Additional inflammatory genes whose differential expression was not deemed statistically significant in our RNA-seq data, e.g., *Cxcl1* and *Il1b*, were in fact downregulated in *Neat1* gRNA iBMDMs (**Fig. S6B-C**). Failure to induce *Rsad2* transcript expression resulted in significantly lower Viperin (RSAD2) protein levels in *Neat1* gRNA iBMDMs compared to off-target gRNA controls (**Fig. 6F**). Based on these data and our BMDM transcriptomics (**Fig. 5**), we hypothesized that *Neat1* gRNA iBMDMs may be compromised in their ability to restrain viral replication. To test this, we infected *Neat1* gRNA and off-target control iBMDMs with vesicular stomatitis virus (VSV) at an MOI of 1. VSV is a single positive strand virus whose replication is exquisitely sensitive to type I interferon and interferon stimulated gene (ISG) expression. We detected a dramatic increase in VSV replication in *Neat1* gRNA iBMDMs at early timepoints postinfection, as measured by RT-qPCR of two VSV genes (*Vsvg* and *Vsvm*) (**Fig. 6H**). Failure to control VSV replication in *Neat1* gRNA iBMDMs was concomitant with low expression of antiviral genes like *Rsad2*, *Mx1*, and *Oas2* (**Fig. 6I**). These data support an important role for *Neat1* in promoting antiviral responses in macrophages.

As M2-like macrophages are known to provide a niche that supports replication of the intracellular bacterial pathogen *Salmonella enterica* serovar Typhimurium (49–51), we hypothesized that *Neat1* gRNA iBMDMs may also be permissive to *Salmonella*. To test this, we infected *Neat1* gRNA and off-target control iBMDMs with *Salmonella* (MOI = 10) and enumerated CFUs from cells at 2 and 20 hrs post infection. We observed a 2-3 fold increase in *Salmonella* replication in iBMDMs lacking *Neat1* (**Fig. 6J**). Immunofluorescence microscopy using an anti-LPS antibody confirmed significantly enhanced *Salmonella* replication in *Neat1* gRNA iBMDMs, without a defect in early internalization (**Fig. 6K and Fig. S6D**). While the precise mechanism through which *Salmonella* can survive and replicate in *Neat1* gRNA iBMDMs remains to be elucidated, these phenotypes are consistent with dysregulated expression of M1/M2 genes like nitric oxide synthase (*Nos2*), a key antimicrobial mediator in macrophages Vazquez-Torres (52, 53), and arginase, *Arg1*, which has been shown to negatively regulate nitric oxide (54, 55) (**Fig. 6L-M**).

## DISCUSSION

Despite enthusiasm surrounding the phenomenon of liquid-liquid phase separation and the structure of MLOs, the function of condensates in cellular homeostasis and stress responses remains poorly understood. Our current knowledge of paraspeckle structure and assembly far exceeds that of paraspeckle function. Here, we investigated a role for paraspeckles in activating the macrophage innate immune response following an infection-relevant stimulus (e.g. LPS). We report that paraspeckles rapidly aggregate, disassemble, and then re-form at steady state levels in a regulated fashion over a very short time-course (approximately 4 hours) in response to multiple innate agonists (**Fig. 1**). During this same time-course, we observed that *Neat1* is needed to induce upregulated inflammatory/antimicrobial genes and repress downregulated wound-healing/cell proliferation-associated genes, which contributes to macrophage control of bacterial and viral replication (**Fig. 5-6**). These findings further our understanding of how biomolecular condensate assembly can be regulated by environmental cues and how dynamic control of these complexes enables rapid calibration of gene expression in the nucleus.

Our experiments identified several unique aspects of the paraspeckle lifecycle in macrophages. First, our data clearly demonstrate that maintenance and upregulation of paraspeckles in activated macrophages requires transcription (**Fig. 3**). As loading of paraspeckles proteins on *Neat1* has been shown to occur co-transcriptionally (31), we know that transcription of *Neat1* itself is needed for PS maintenance. However, whether *Neat1* itself is transcriptionally <u>up</u>regulated as part of the response to LPS remains unclear. By RT-qPCR, we typically measure a 2- to 3-fold increase in *Neat1* transcript levels at 1 hr post-LPS treatment (**Fig. 1B**). This peak in total transcript is later than the peak of PS aggregation, which occurs at 0.5 hr, suggesting that early PS aggregation is driven by sequestration of already synthesized *Neat1* as opposed to *de novo* transcription of new *Neat1* transcripts. This makes sense, given the rapid aggregation of PS observed (∼0.5 hr) and the length of *Neat1* itself (21.1 kb), which will take some time to fully transcribe. Previously published RNA sequencing data and ChIP-seq data for the NFκB transcription factor subunit RelA (56) show some evidence for RelA binding at the *Neat1* promoter following LPS stimulation of BMDMs (**Fig. S3D**), although this was not concomitant with significantly increased *Neat1* sequencing reads (**Fig. S3E**). These data and our findings together support a model whereby paraspeckle aggregation in macrophages does not require cells to make more *Neat1*. With that said, it remains a possibility that bulk measurements of *Neat1* RNA from a population of cells obscures our ability to accurately measure *Neat1* transcriptional activation. Promoter fusion constructs and/or single cell transcriptomics may help answer the question of whether *Neat1* is transcriptionally induced downstream of pattern recognition receptor engagement and help reconcile our findings with other studies whose models invoke *Neat1* transcriptional upregulation following PAMP sensing in non-immune cells (28).

We can also conclude that the RNA exosome, specifically the NEXT targeting complex, is involved in turning over *Neat1* and regulating paraspeckle dynamics in macrophages (**Fig. 4**). This is consistent with another report linking MTR4 and NEXT to *Neat1* turnover, although the kinetics of *Neat1* turnover reported in HeLa cells (loss of detectable *Neat1* transcript by 6-8 hours post-transcription shut-off) are different from what we see in macrophages (2 hours) (57). The fact that we see paraspeckle upregulation in resting *Mtr4*, *Zcchc8*, and *Dis3* knockdown macrophages demonstrates a role for the exosome in constitutively controlling paraspeckle size/numbers. The fact that we see paraspeckles maintained in exosome-knockdown macrophages even at timepoints when paraspeckles disaggregate in wild-type cells (2 hrs post-LPS) can mean two things: 1.) exosome targeting of *Neat1* is enhanced at 2 hrs post-LPS treatment, suggesting that the exosome itself is regulated downstream of pattern recognition receptor engagement or 2.) exosome turnover of *Neat1* is constant and *Neat1* transcription is turned off in a regulated fashion. Potential mechanisms for such a turn off could include competition between the long and short isoforms of *Neat1*, histone modification/chromatin remodeling at the *Neat1* promoter, and altered association of the *Neat1* genomic locus with enhancer elements. Again, promoter fusions and/or cell lines that enable rapid loss of exosome components (like the auxin degron system in (58)) could help tease apart the role of transcription versus turnover in regulating *Neat1* levels post-LPS treatment.

One curious observation made regarding paraspeckles in macrophages is their disintegration at 2 hrs post-LPS treatment. To the best of our knowledge, this is the first report of paraspeckle ablation following a physiologically-relevant cellular stress. At 2 hours post-LPS treatment, the macrophage innate response is still ramping up, with maximum transcript accumulation for many inflammatory and antimicrobial genes seen at 4-6 hrs post-stimulation. Break-down of paraspeckles at 2 hours post-LPS could aid in this amplification by, for example, releasing factors involved in nucleosome remodeling at secondary response genes (e.g., SWI/SNF components (36)) or post-transcriptional processing of innate transcripts (e.g., RBPs). Such a model would make paraspeckle disintegration a critical step in activation of inflammatory responses and may position the paraspeckle as a stopgap to prevent this “ramping up” (i.e., continue to sequester innate activating proteins) in the absence of a strong stimulus. On the other hand, it is possible that early paraspeckle aggregation sequesters inhibitory factors to help activate the innate response. Such a model has been proposed for paraspeckle-mediated relief of SFPQ repression of *IL8* transcription in HeLa cells (28). Consistent with this idea, we observed increased colocalization between the repressive RNA binding protein hnRNP M and *Neat1* following LPS treatment (**Fig. 2D**). Our previous work showed that hnRNP M can repress innate gene expression in macrophages by slowing intron removal in inflammatory transcripts like *Il6* (38). Thus, is it possible that dampened IL-6 expression in *Neat1* KO macrophages (**Fig. 5G-I** and **6D**) is driven in part by failure to sequester hnRNP M from nascent pre-mRNAs. HnRNP M is likely one of several RBPs whose paraspeckle association is stimulated by macrophage activation. The dual action of *Neat1* in promoting expression of genes turned on by LPS and dampening expression of genes turned off by LPS suggests that paraspeckles can sequester RBPs with opposing functions to simultaneously activate and inhibit expression of innate genes. While we do not fully understand the molecular mechanisms driving these phenomena, we can conclude that the nuclear paraspeckle plays a crucial and underappreciated role in globally regulating the macrophage innate response to pathogens.

## ACKNOWLEDGEMENTS

The authors would like to thank members of the Patrick and Watson labs for helpful discussions and manuscript edits. Our sincerest appreciation goes to Malea Murphy and the Integrated Microscopy and Imaging Laboratory (IMIL) at Texas A&M University School of Medicine, who helped with acquisition and analysis of our FISH-IF images. We would also like to acknowledge the Genomics and RNA Profiling Core at Baylor College of Medicine for library generation and high-throughput sequencing and the CRISPR Core at the University of California, Santa Cruz (RRID:SCR_021207) for generating gRNA-expressing lentivirus.

## MATERIALS AND METHODS

### RNA FISH and immunofluorescence microscopy

Fluorescence In Situ Hybridization-Immunofluorescence (FISH-IF) was used to simultaneously visualize the *Neat1* lncRNA and proteins. Briefly, 1 x 10^5^ RAW 264.7 macrophages or BMDMS were plated in a glass bottom 35mm Mattek dish and allowed to rest for 24h. Cells were washed with 1xPBS and fixed with 4% paraformaldehyde for 10 min, and then washed 3x with 1xPBS. Cells were permeabilized with 0.5% Triton X-100 for 10 min, washed twice with PBS and once with 2xSSC buffer (Sigma, S6639) for 10 min. Cells were incubated overnight with probes in hybridization buffer (1:200). A pool of 48 *Neat1* smFISH (single molecule Fluorescence In-Situ Hybridization-Immunofluorescence) RNA probes were designed and purchased from Stellaris® (SMF-3010-1, Mouse *Neat1* Middle Segment with Quasar® 570 Dye). Following probe incubation, cells were washed with 2xSSC each time for 15 min, and then 3 times with PBS. DAPI (Invitrogen, D1306, 1:10,000) was used for 10 min for nuclear staining, followed by four PBS washes for 5 min each. For immunofluorescence, primary antibodies (1:200) against SFPQ (Abcam, ab177149), PSP1 (Abcam, ab214012), NONO (Abcam, ab133574), HNRNP M (Abcam, ab177957), FUS (Abcam, ab243880), BRM (Abcam, ab240648), BRG1 (Abcam, ab110641), NF-κB (Active Motif, Catalog No: 40916) were used along with 10% BSA (NEB #B9200) as a blocking agent. Cells were overnight incubated with primary antibody at 4°C, and then washed with PBS three times for 5 min each. Cells were then stained with fluorescent secondary antibodies (goat anti-rabbit Alexa Fluor 488, and/or goat anti-mouse Alexa Fluor 488; Invitrogen, 1:1,000), and finally washed with PBS three times for 5 min each. Images were taken on an Olympus Fluoview FV3000 confocal laser scanning microscope using the 60X objective and processed and analyzed with Fiji.

### Primary Cell Culture

To prepare primary cell cultures of bone marrow-derived macrophages (BMDMs), bone marrow (BM) cells were isolated from mouse femurs by washing them with 10 mL DMEM 1 mM sodium pyruvate (ThermoFisher, 11995065), followed by centrifugation for 5 min at 400 RCF and resuspension in BMDM media consisting of Dulbecco modified Eagle medium (DMEM), 20% of heat-inactivated fetal bovine serum (FBS) (Millipore, F0926), 1 mM sodium pyruvate (Lonza, BE13-115E), and 10% MCSF conditioned media. BM cells were counted and seeded at a density of 5x10^6^ cells per 15 cm non tissue culture-treated dish in 30 ml complete BMDM media. An additional 15 mL of BMDM media was added on day 3, and cells were harvested on day 7 using 1 X PBS EDTA (Lonza, BE02-017F) for experiments.

### Cell Lines and Cell Culture

Low passage stocks of RAW 264.7 macrophages were obtained from ATCC (TIB-71), were cultured at 37°C/5% CO2 in complete media containing high glucose DMEM (Thermo Fisher, 11965092), with 10% FBS (Millipore, F0926) and 0.2% HEPES (Thermo Fisher, 15630080). Cells were harvested using 1xPBS + EDTA or cell scraper. Absence of mycoplasma was confirmed in all cell lines using the Universal Mycoplasma Detection Kit (ATTC, 30–1012 K). dCas9 iBMDMs obtained from the Carpenter Lab at University of California, Santa Cruz (UCSC), were cultured at 37°C/5% CO_2_ in complete media containing high glucose DMEM (Thermo Fisher, 11965092), with 10% FBS (Millipore, F0926) and 0.2% HEPES (Thermo Fisher, 15630080). To make *Neat1* gRNA-expressing cell lines, 2.5 x 10^4^ cells were plated in a 24-well treated tissue culture plate and incubated for 24 hrs. The cells were then transduced with *Neat1* gRNA or off-target gRNA lentivirus generated by the UCSC CRISPR Core in the presence of 0.5 μl lipofectamine 2000 (Thermo Fisher, 52887). Twenty-four hours after initial transduction, lentivirus was removed and replaced with complete media containing high glucose DMEM for another 24 hrs. 48h post-transduction, cells were selected via the addition of 3μg/mL puromycin (Invivogen, ant-pr-1). 96 hrs post-transduction, cells were selected in final concentration of 5μg/mL puromycin and maintained in this concentration of drug.

### Cell Stimulations and Treatments

BMDMs/RAW 264.7 macrophages were plated at 5 x 10^5^ cells/well in 12-well dishes. For FISH and IF-FISH, 1 x 10^5^ cells were plated into 35mm Mattek plates a day prior to stimulation. LPS stimulations were done with 100 ng/ml LPS (InvivoGen) for RAW 264.7 macrophages and 10ng/ml LPS for BMDMs for times indicated. Cells were treated with 100 ng/ml Pam3CSK4 (InvivoGen) or transfected with 1 μg/ml ISD, or 500 ng/ml poly(I:C) using lipofectamine (Thermo Fisher). For transcription inhibition, 5x10^5^ cells/well were plated in a 12-well dish or 35mm Mattek plates and transcription was blocked by using 5 μg/ml actinomycin D for 30min. For kinase inhibitor experiments, cells were plated in 35mm Mattek/12-well plates and treated with P38 inhibitor (SB203580) 10 μM, MEK1/2 Inhibitor (U0126) 25 μM or JNK inhibitor (SP600125) 25 μM, (InvivoGen) for 45 min prior to imaging.

To polarize RAW 264.7 macrophages into M1/M2 macrophages, RAW 264.7 macrophages were plated in a glass bottom 35mm Mattek (P35G-1.5-14-C) or 12 well plates and treated with 50 ng/mL of IFN-γ (R & D, 485-MI) or 25 ng/mL of IL-4 (PeproTech, 214-14) for 24 hours. Macrophages polarization was confirmed by gene expression analysis.

### *Samonella* infections

*Salmonella enterica* serovar Typhimurium (SL1344) was obtained as a kind gift from Dr. Denise Monack at Stanford University. A bacterial inoculum was prepared by growing the bacteria to the log growth phase with an optical density at 600nm (OD600) of 1.0. The bacterial culture was then centrifuged at 500 rpm for 5 min to remove the LB media. The bacteria were subsequently washed with Hank’s Balanced Salt Solution (HBSS) by pelleting at 3000 rpm for 5 min. Bacteria were suspended in HBSS and added to the macrophages, followed by a brief centrifugation step. The cells were then incubated at 37°C for 20 min. After the incubation period, the cells were washed three times with HBSS and fresh media was added for the specified time courses. Prior to cell lysis, the cell culture media was aspirated, and the cells were washed twice with phosphate-buffered saline (PBS). For cell lysis, 1mL of cell lysis buffer (PBS containing 1% Triton X-100 and 0.1% SDS) was added directly to each well. The lysate was carefully transferred to an Eppendorf tube, followed by vortexing for 5-10 seconds to ensure thorough mixing. To prepare serial dilutions, 10-fold dilutions were performed up to 10^-4 using PBS. The plates were then incubated overnight at 37°C. Colony-forming units (CFUs) were counted, and the CFU/mL and fold replication were determined using the following calculations:

CFU/mL = (number of colonies / dilution factor) × volume plated.

### Vesicular stomatitis viral (VSV) infections

*Neat1* gRNA or off-target gRNA iBMDMs were seeded in 12-well tissue culture-treated plates at a density of 7×10^5^ cells per well 12h before infection. The next day cells were infected with VSV-GFP virus at multiplicity of infection (MOI) of 1 in serum-free DMEM (DMEM Thermo Fisher, 11965092). After 1hr of incubation with media containing virus, supernatant was removed, and fresh DMEM plus 10% FBS (Millipore, F0926) was added to each well. At indicated time points post infection, cells were harvested with TRIzol for RNA isolation.

### siRNA Transfections

To perform mRNA knockdown of exosome components, cells were plated in 12-well plates at a density of 3 x 10^5^ RAW 264.7 macrophages or 3.5 x 10^5^ BMDMs on day 4 of differentiation and rested overnight. The next day, complete media was replaced with 500 μL fresh complete media 30 min prior to transfection. Transfection was carried out using Fugene SI reagent (SKU:SI-100) along with 50μM of ThermoFisher siRNA stock against Skiv2l2 (Mtr4); s90745), Zcchc8; s89034, Zcchc7; s38623, Dis3; s91196, and Exosc10; s78572). For a negative control, Silencer Select Negative Control #1 (ThermoFisher, 4390843) was used. Cells were incubated for 48h in transfection media at 37°C with 5% CO_2_ before conducting downstream experiments.

### *Neat1* KO Mice genotyping

*Neat1* KO mice used in this study were obtained from the laboratory of Shinichi Nakagawa at Hokkaido University (46). Briefly, a BAC clone RP23-209P9 was used as a template to amplify DNA fragments through PCR, which were subsequently subcloned into DT-ApA/LacZ/NeO to create the targeting vector. Homologous recombination was verified through Southern blot analysis after electroporation of the linearized targeting vector into TT2 embryonic stem cells. Chimeric mice were produced using the recombinant embryonic stem clone and crossed with C57BL/6 females to generate NEAT1lacZ/Neo/+ heterozygous animals. These mice were then bred with Gt (ROSA) 26 Sortm1 (FLP1) Dym (The Jackson Laboratory) to flip out the PGK-Neo cassette. The resulting heterozygous mice (NEAT1 lacZ/+) were maintained on the C57BL/6 genetic background and genotyped through PCR using DNA obtained from ear clips. The genotyping primer sequences are NEAT1 WT FW: CTAGTGGTGGGGAGGCAGT, NEAT1 WT RV: AGCAGGGATAGCCTGGTCTT, LacZ 5(KO) RV: GCCATTCAGGCTGCGCAACTG. The mice used in our experiments were age- and sex-matched controls, with 8-12 week old mice used to generate BMDMs. All mice were housed, bred, and studied at Texas A&M Health Science Center in accordance with approved Institutional Care and Use Committee guidelines.

### RNA Sequencing

RNA from *Neat1^-/-^* and WT BMDMs was isolated using TRIzol. Ribodepletion library preparation was carried out by the Baylor College of Medicine Genomic and RNA Profiling Core (GARP) in biological triplicate. RNA sequencing (150bp paired-end reads) was performed on an Illumina NovaSeq 6000 with S4 flow cell. STAR alignment to the *Mus musculus* Reference genome (mm10) and differential expression analysis was carried out using Illumina BaseSpace RNA-Seq Alignment and RNA-seq Differential Expression apps. Differentially expressed genes were identified based on an adj. p-value threshold of <0.05 and a log_2_FC of +/- 0.5. For transcriptome analysis, Qiagen IPA analysis was utilized to generate lists of GO terms and disease pathways.

### Immunoblot

Cells were washed with PBS and lysed in 1X RIPA buffer with protease and Pierce EDTA free phosphatase inhibitors (Thermo Scientific, A32957) with 1 U/mL of benzonase nuclease (Millipore, 101697) added to degrade genomic DNA. Proteins were separated by SDS-PAGE on Any kD mini-PROTEAN TGX precast gel (BioRad) and transferred to 0.45 μm nitrocellulose membranes (Cytiva, 10600041). After blocking the membranes for 1 hour at RT in LiCOR Odyssey blocking buffer (927–60001), they were incubated overnight at 4°C with the relevant primary antibodies, including β-ACTIN (Abcam 6276, 1:5000), SFPQ (ab177149, 1:1000), PSP1 (ab214012, 1:1000), NONO (ab133574, 1:1000), Anti-phosphoepitope SR proteins, clone 1H4 (EMD Millipore, MABE50, 1:1000). Membranes were washed three times for 5 min in PBS-Tween-20 and incubated with the appropriate secondary antibodies (LI-COR, 925-32210, 926-68071) for 1 hour at RT before imaging on a LiCOR Odyssey Fc Dual-Mode Imaging System.

### RNA Isolation and RT-qPCR Analysis

RNA was isolated from cells harvested in TRIzol using Direct-zol RNA Miniprep kits (Zymo Research, R2052) with an hour DNase treatment. cDNA was synthesized using iScript cDNA Synthesis Kit (Bio-Rad, 1708891) and diluted to 1:20 for each sample. A standard curve was generated using a pool of cDNA from each treated sample. RT-qPCR was performed using Power-Up SYBR Green Master Mix (Thermo Fisher, A25742) and a QuantStudio Flex6 (Applied Biosystems) with triplicate wells in a 384-well plate.

### Data Analysis and Presentation

Graphpad Prism software (Version 9) was used to perform statistical analysis of data and generate graphs. Unless otherwise indicated, the results presented are from at least three biological samples and are represented as the mean, with error bars representing SEM. We used a t test and one/two-way analysis of variance (ANOVA) to determine significant differences between the means of the control groups versus experimental groups. Quantification of PS area was carried out by measuring the 3D area of signal in ≥100 cells using Fiji software. RNA and protein colocalization was analyzed using the Coloc 2 plugin in Fiji (ImageJ) software. Briefly, the images were first background-subtracted and thresholded to generate binary masks of the *Neat1* RNA and protein signals. The Pearson correlation coefficient, a quantitative measure of the degree of co-localization between the two signals, was then calculated and graphed.

### Primers

Sequences of the primer sets used in this paper are as follows:

Neat1-V2-Fwd: GTGGTCTCTGTGGAAGTGTATG

Neat1-V2-Rev: TGGAGAAGCGAAACGAGATG,

TNF-Fwd: CCGATGGGTTGTACCTTGTC

TNF-Rev: AGATAGCAAATCGGCTGACG

IL-1B-Fwd: GGTGTGTGACGTTCCCATTA

IL-1B-Rev: ATTGAGGTGGAGAGCTTTCAG

Malat1-Fwd: GATGACTCAAGGGAACCAGAAA

Malat1-Rev: GAAAGCTAGCATCCATCCTCT

AC Nos2-Fwd: GCAGCACTTGGATCAGGAA

Nos2-Rev: GAAACTTCGGAAGGGAGCAA

GAPDH-Fwd: CAATGTGTCCGTCGTGGATCT

GAPDH -Rev: GTCCTCAGTGTAGCCCAA GAT

Kcnq1ot1-Fwd: CGTTATCCAGACTCTCCCTTTC

Kcnq1ot1-Rev: GTTACCTCTTTCAGGGCT

TCT MTR4-Fwd: CCCACTCCACAATGATCCTAAC

MTR4-Rev: CTTGCCTTCTTCAGTTCTCTCTT

Dis3-Fwd: ACACACATTTCACCTCTCCTATC

Dis3-Rev: GTCAACTCAGGGTAAGTACAGTC

Exosc10-Fwd: CTGTTTGCTTGGAGGGATAAGA

Exosc10-Rev: GGCAGTTCCTCAGCTATCTTTAG

Zcchc7-Fwd: GGAGGAGAGGACATAGCAAATAC

Zcchc7-Rev: CAAGTTTGAGGATGCCTTTCAC

ZCCHC8-Fwd: GCTTACGGAAGGATGGGAAATA

ZCCHC8-Rev: CAGTTGAAACAGTGAGGCTTTG

---

**Figure S1:**
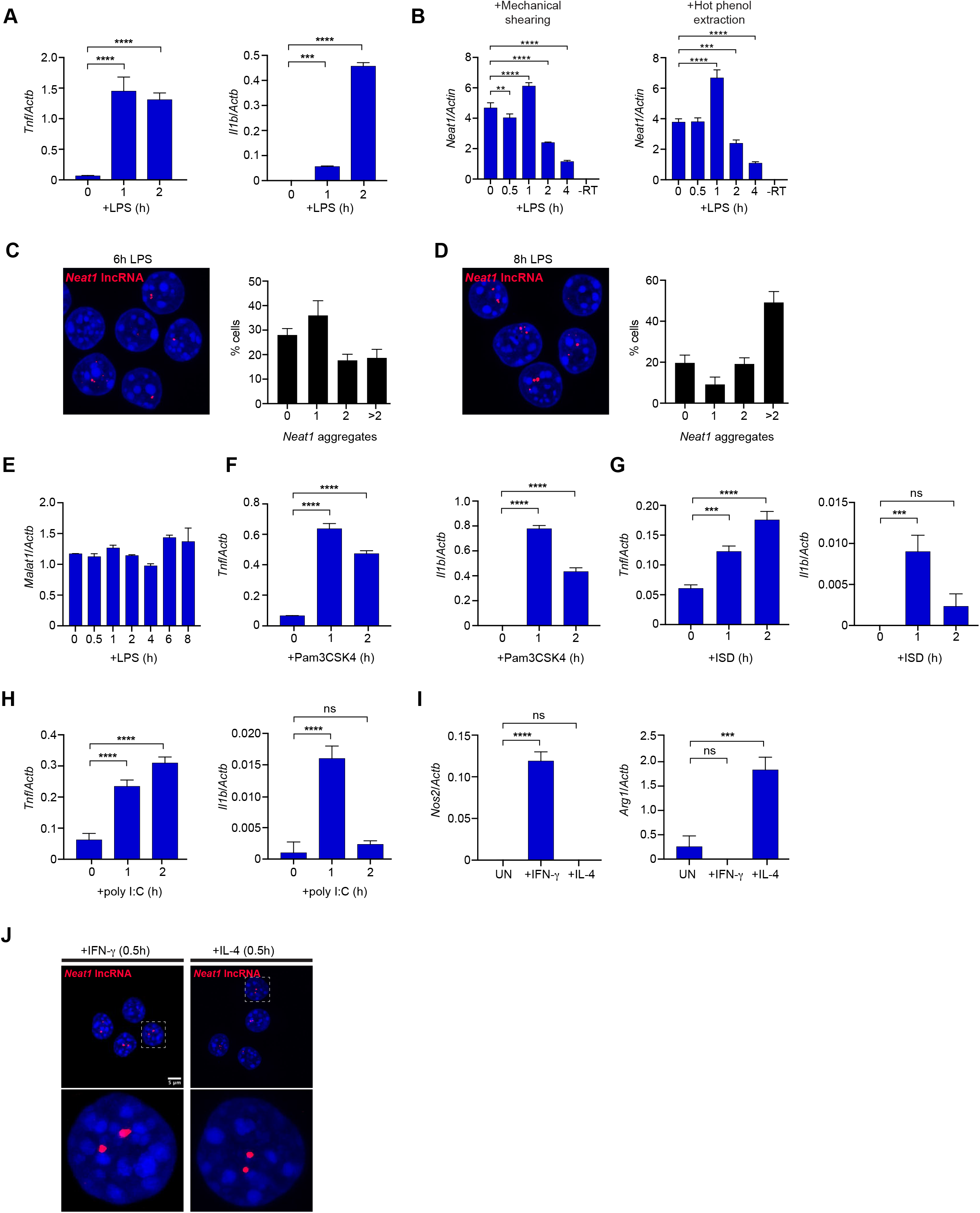
Supplementary Data for Figure 1. (A) RT-qPCR of *Tnfa* and *Il1b* transcript levels in RAW 264.7 macrophages post-LPS treatment, (100ng/ml) shown relative to *Actb.* (B) RT-qPCR of Neat1 in RAW 264.7 macrophages over a 4h LPS treatment after three different RNA extraction methods: (left) mechanical shearing (100 passages through 20-gauge needle), or (right) hot phenol extraction (55 deg for 10 min with 250 rpm shaking). (C) RNA-FISH of *Neat1* (red) in RAW 264.7 macrophages at 6 hrs post-LPS (100 ng/ml) treatment. To quantify on right, cells with different numbers of *Neat1* aggregates were manually counted (n>100) and binned into cells with 0, 1, 2, and >2 paraspeckles. (D) As in (C) but at 8 hrs post-LPS stimulation. (E) RT-qPCR of *Malat1* transcript levels in RAW 264.7 macrophages post-LPS treatment (100ng/ml) shown relative to *Actb.* (F) As in (A) but post-Pam3CSK4 (100 ng/ml) treatment. (G) As in (A) but post-dsDNA (ISD) (1 mg/ml) transfection. (H) As in (A) but post-dsRNA (polyI:C) (500 ng/ml) transfection. (I) RT-qPCR of M1-(*Nos2*) and M2-characteristic (*Arg1*) transcripts 24 hrs post-IFN-γ (50 ng/ml) or IL-4 (25 ng/ml) treatment, shown relative to *Actb.* (J) RNA-FISH of *Neat1* (red) in RAW 264.7 macrophages at 0.5 hr post-IFN-γ (50 ng/ml) or IL-4 (25 ng/ml) treatment. <u>Statistical tests</u>: Data is presented as the mean of three biological replicates unless otherwise noted with error bars representing SEM. At least 100 cells were counted over multiple coverslips per condition. Statistical significance was determined using a one-way ANOVA (A-I). *p<0.05, **p<0.01, ***p<0.001, ****p<0.0001.

**Figure S2:**
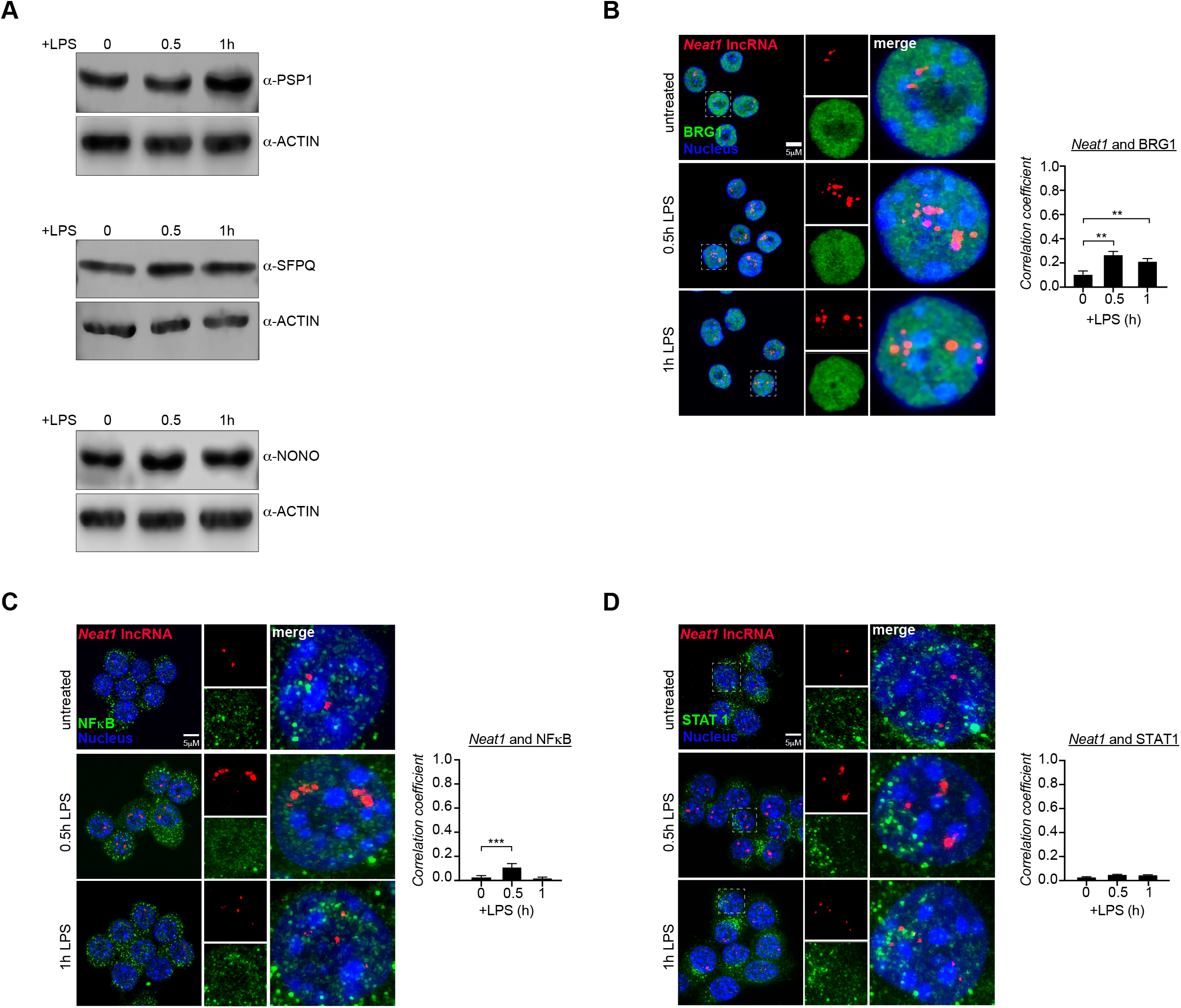
Supplementary Data for Figure 2. (A) Immunoblot analysis of PSP1, SFPQ, and NONO in RAW 264.7 macrophages post-LPS (100 ng/ml) stimulation. ACTB used as a loading control. Representative blot of n=3. (B) RNA-FISH of *Neat1* (red) and immunofluorescence microscopy of BRG1 (green) in RAW 264.7 macrophages at 0.5 and 1 hr post-LPS treatment (100 ng/ml). Correlation coefficient between *Neat1* and BRG1 quantified to the right. (C) As in (B) but for NFκB (RelA/p65). (D) As in (B) but for STAT1. <u>Statistical</u> <u>tests</u>: Data is presented as the mean of three biological replicates unless otherwise noted with error bars representing SEM. At least 100 cells were counted over multiple coverslips per condition. Colocalization coefficient was measured using the ImageJ plugin Coloc2. Statistical significance was determined using a one-way ANOVA (B-D). **p<0.01, ***p<0.001.

**Figure S3:**
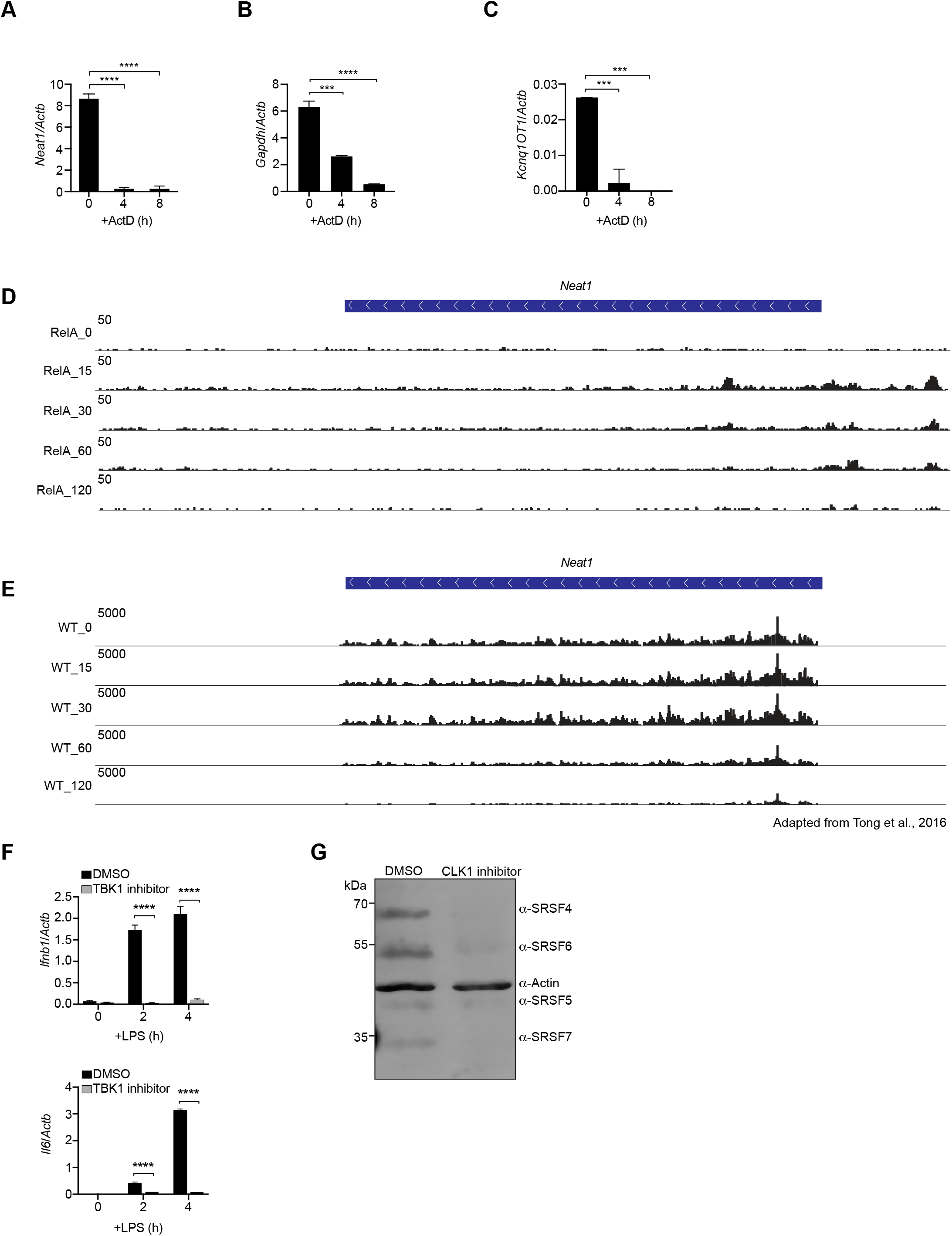
Supplementary Data for Figure 3. (A) RT-qPCR of *Neat1_2* transcript levels in RAW 264.7 macrophages after 4 and 8 hrs ActD treatment, (5mg/ml) shown relative to *Actb*. (B) As in (B) but for *Gapdh*. (C) As in (B) but for *Kcnq1OT1*. (D) IGV tracks of ChIP-seq data for RelA (NFκB subunit) at the *Neat1* promoter over a 2 hr time course of LPS treatment. From Tong et al., 2016. GEO Accession numbers: GSM1645112 (RelA_0), GSM1645114 (RelA_15), GSM1645116 (RelA_30), GSM1645118 (RelA_60), GSM1645120 (RelA_120). (E) IGV tracks of RNA-seq reads at *Neat1* over a 2 hr time course of LPS treatment. From Tong et al., 2016. GEO Accession numbers: GSM1645338 (WT_0), GSM1645340 (WT_15), GSM1645342 (WT_30), GSM1645344 (WT_60), GSM16453346 (WT_120). (F) RT-qPCR of *Ifnb1* and *Il6* transcript levels in LPS-treated RAW 264.7 macrophages +/- the TBK1 inhibitor (GSK-8612; 10 mM). (G) Immunoblot analysis of phosphorylated SR proteins in RAW 264.7 macrophages +/- treatment with the CLK1 inhibitor (Cpd 23; 10 mM). <u>Statistical tests</u>: Data is presented as the mean of three biological replicates unless otherwise noted with error bars representing SEM. Statistical significance was determined using a one-way ANOVA (A-C) or Student’s t-test (F) or. ***p<0.001, ****p<0.0001.

**Figure S4:**
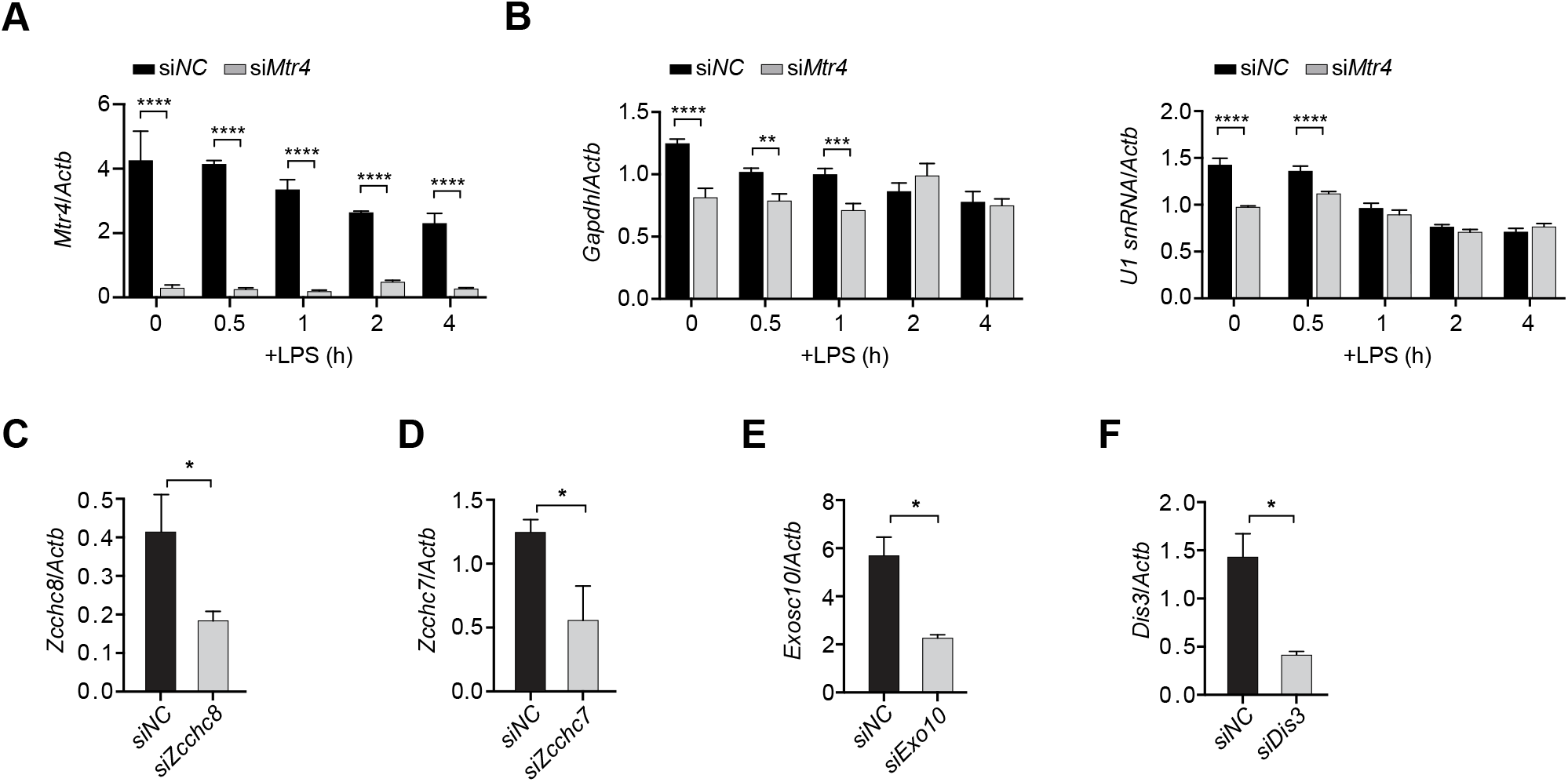
Supplementary Data for Figure 4. (A) RT-qPCR of *Mtr4* transcript levels in *siNC* and *siMtr4* RAW 264.7 macrophages over a timecourse of LPS treatment (100 ng/ml), shown relative to *Actb*. (B) As in (A) but for *Gapdh* and *U1 snRNA,* shown relative to *Actb*. (C) RT-qPCR of *Zcchc8* transcript levels in *siNC* and *siZcchc8* RAW 264.7 macrophages, shown relative to *Actb*. (D) As in (C) but for *Zcchc7* in *siNC* and *siZ-cchc7* RAW 264.7 macrophages. (E) As in (C) but for *Exosc10* in *siNC* and *siExosc10* RAW 264.7 macrophages. (F) As in (C) but for *Dis3* in *siNC* and *siDis3* RAW 264.7 macrophages. <u>Statistical</u> <u>tests</u>: Data is presented as the mean of three biological replicates unless otherwise noted with error bars representing SEM. Statistical significance was determined using a Student’s t-test (A-F). *p<0.05, **p<0.01, ***p<0.001, ****p<0.0001.

**Figure S5:**
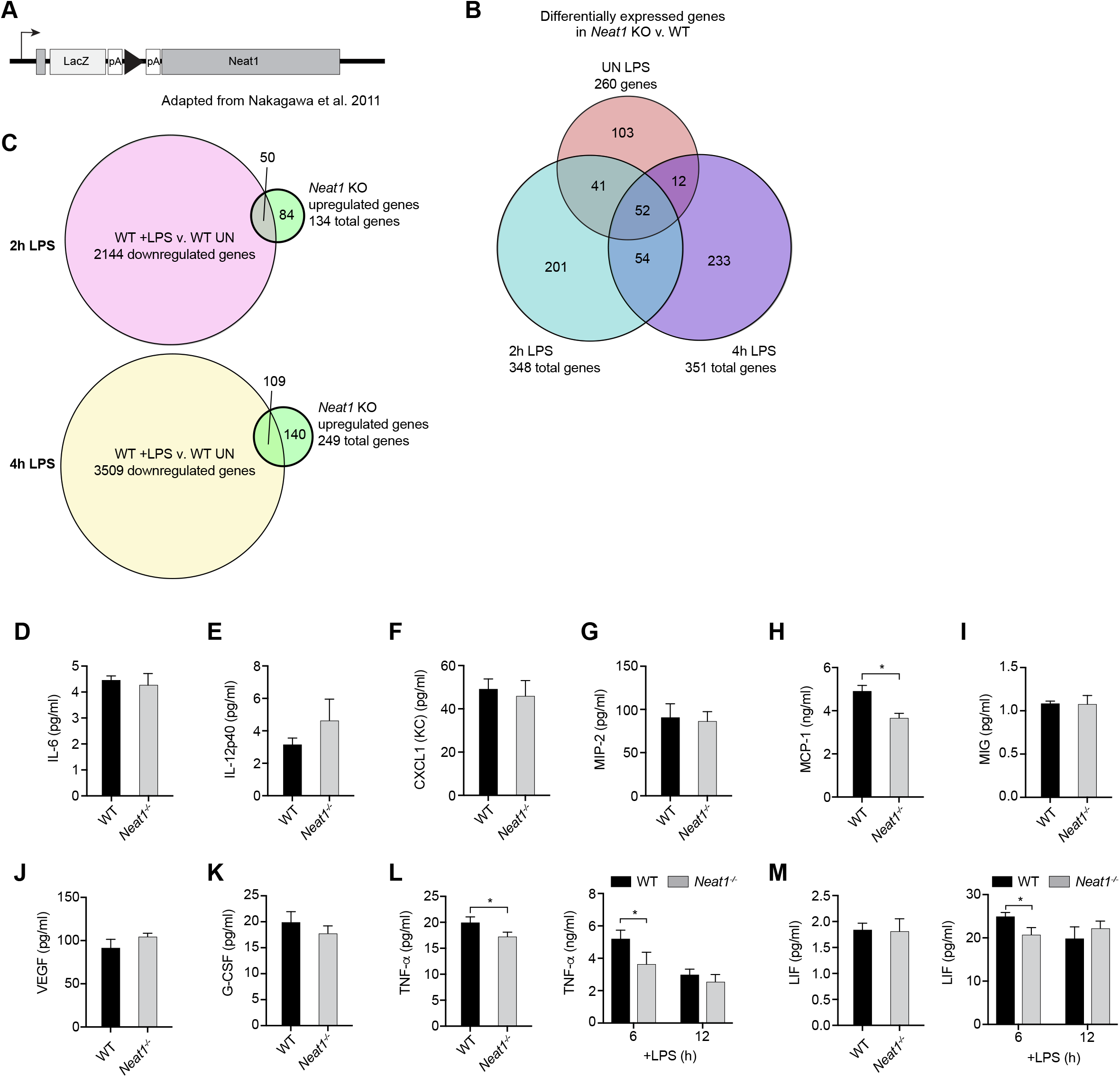
Supplementary Data for Figure 5. (A) Diagram of Neat1/lacZ (*Neat1* KO) mice. (B) Venn diagram of differentially expressed genes in *Neat1* KO v. WT BMDMs in untreated, 2 hrs post-LPS, and 4 hrs post-LPS. (C) (top) Venn diagram of genes downregulated as part of the WT BMDM response to LPS (WT 4 hrs post-LPS v. WT UN; yellow) compared to genes upregulated at 4 hrs post-LPS in *Neat1* KO BMDMs (green). (bottom) Venn diagram of genes downregulated as part of the WT BMDM response to LPS (WT 2 hrs post-LPS v. WT UN; pink) compared to genes upregulated at 2 hrs post-LPS in *Neat1* KO BMDMs (green). (D) Measurements of IL-6 levels in the supernatants of WT and *Neat1* KO BMDM supernatants at rest via cytokine array. (E) As in (D) but for IL-12p40. (F) As in (D) but for KC (CXLC1). (G) As in (D) but for MIP-2 (CXCL2). (H) As in (D) but for MCP-1 (CCL2). (I) As in (D) but for MIG (CXCL9). (J) As in (D) but for VEGF. (K) As in (D) but for G-CSF. (L) Measurements of TNF levels in the supernatants of WT and *Neat1* KO BMDM supernatants at rest and at 6 and 12 hrs post-LPS treatment via cytokine array. (M) As in (L) but for LIF. <u>Statistical tests</u>: Data is presented as the mean of three biological replicates unless otherwise noted with error bars representing SEM. Statistical significance was determined using a Student’s t-test (D-M). *p<0.05, **p<0.01, ***p<0.001, ****p<0.0001.

**Figure S6:**
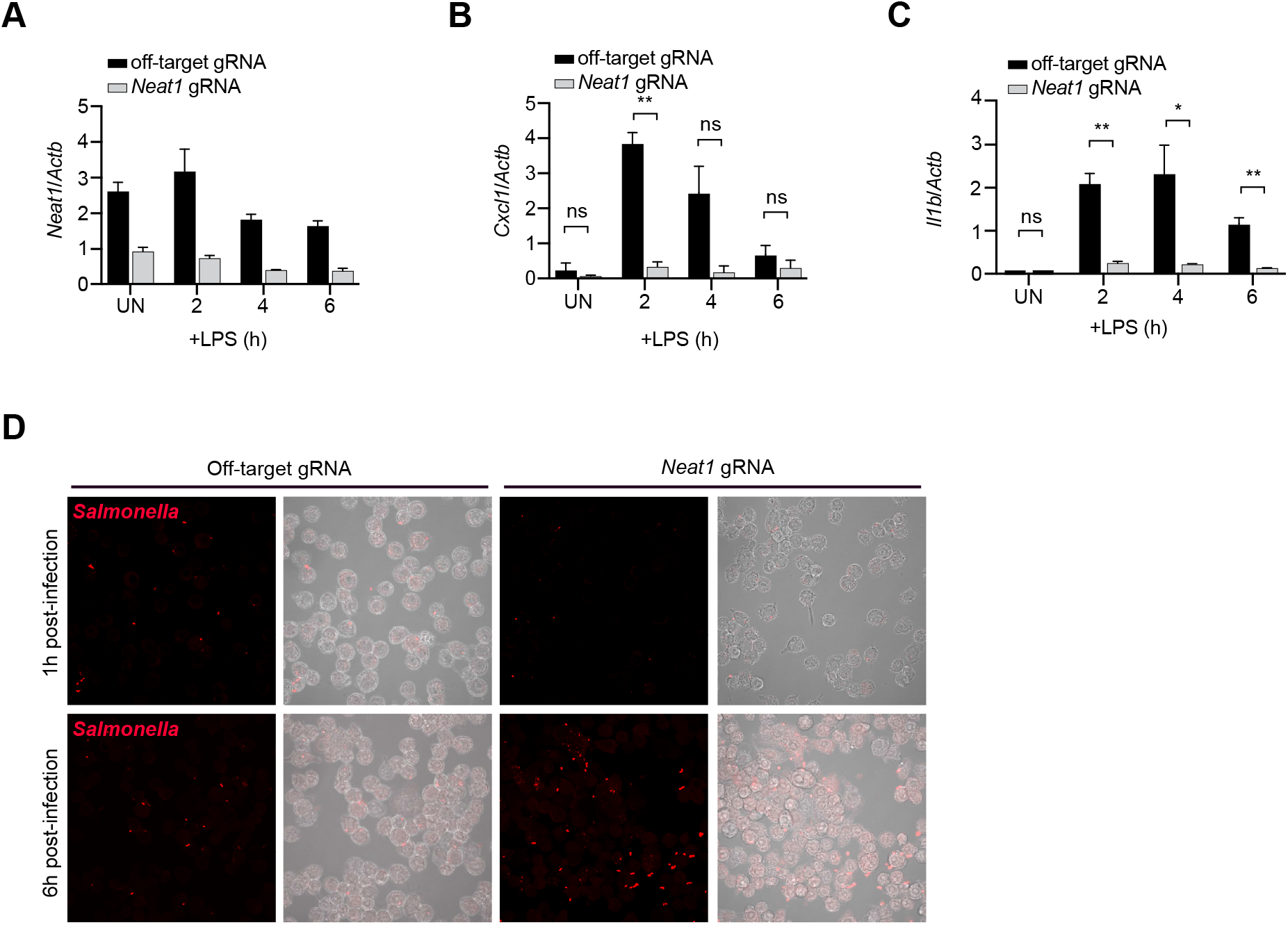
Supplementary Data for Figure 6. (A) RT-qPCR of *Neat1_2* transcript levels in off-target gRNA and *Neat1* gRNA-expressing dCas9 iBMDMs after LPS treatment (100 ng/ml) shown relative to *Actb*. (B) RT-qPCR of *Cxcl1* transcript levels in off-target gRNA and *Neat1* gRNA dCas9-expressing iBMDMs at 2, 4, and 6 hrs post-LPS (100 ng/ml), shown relative to *Actb*. (C) As in (B), but for *Il1b*. (D) Immunofluorescence of *Salmonella* (anti-LPS antibody; red) at 1 and 6 hr post-*Salmonella* infection (MOI=10) in off-target gRNA and *Neat1* gRNA dCas9-expressing iBMDMs (multi-cell view). <u>Statistical tests</u>: Data is presented as the mean of three biological replicates unless otherwise noted with error bars representing SEM. Statistical significance was determined using a Student’s t-test (A-C). *p<0.05, **p<0.01, ***p<0.001, ****p<0.0001.

## REFERENCES

1. E. Gomes, J. Shorter, The molecular language of membraneless organelles. J Biol Chem 294, 7115–7127 (2019).

2. W. van Leeuwen, C. Rabouille, Cellular stress leads to the formation of membraneless stress assemblies in eukaryotic cells. Traffic 20, 623–638 (2019).

3. I. A. Sawyer, D. Sturgill, M. Dundr, Membraneless nuclear organelles and the search for phases within phases. Wiley Interdiscip Rev RNA 10, e1514 (2019).

4. E. M. Courchaine, A. Lu, K. M. Neugebauer, Droplet organelles? EMBO J 35, 1603–1612 (2016).

5. H. Ismail et al., Mechanisms and regulation underlying membraneless organelle plasticity control. J Mol Cell Biol 13, 239–258 (2021).

6. C. S. Bond, A. H. Fox, Paraspeckles: nuclear bodies built on long noncoding RNA. J Cell Biol 186, 637–644 (2009).

7. A. H. Fox et al., Paraspeckles: a novel nuclear domain. Curr Biol 12, 13–25 (2002).

8. J. A. West et al., Structural, super-resolution microscopy analysis of paraspeckle nuclear body organization. J Cell Biol 214, 817–830 (2016).

9. A. H. Fox, C. S. Bond, A. I. Lamond, P54nrb forms a heterodimer with PSP1 that localizes to paraspeckles in an RNA-dependent manner. Mol Biol Cell 16, 5304–5315 (2005).

10. A. H. Fox, S. Nakagawa, T. Hirose, C. S. Bond, Paraspeckles: Where Long Noncoding RNA Meets Phase Separation. Trends Biochem Sci 43, 124–135 (2018).

11. T. Yamazaki et al., Functional Domains of NEAT1 Architectural lncRNA Induce Paraspeckle Assembly through Phase Separation. Mol Cell 70, 1038–1053 e1037 (2018).

12. T. Naganuma et al., Alternative 3’-end processing of long noncoding RNA initiates construction of nuclear paraspeckles. EMBO J 31, 4020–4034 (2012).

13. Y. T. Sasaki, T. Ideue, M. Sano, T. Mituyama, T. Hirose, MENepsilon/beta noncoding RNAs are essential for structural integrity of nuclear paraspeckles. Proc Natl Acad Sci U S A 106, 2525–2530 (2009).

14. C. M. Clemson et al., An architectural role for a nuclear noncoding RNA: NEAT1 RNA is essential for the structure of paraspeckles. Mol Cell 33, 717–726 (2009).

15. P. M. Gordon, F. Hamid, E. V. Makeyev, C. Houart, A conserved role for the ALS-linked splicing factor SFPQ in repression of pathogenic cryptic last exons. Nat Commun 12, 1918 (2021).

16. L. V. W. Stagsted, E. T. O’Leary, K. K. Ebbesen, T. B. Hansen, The RNA-binding protein SFPQ preserves long-intron splicing and regulates circRNA biogenesis in mammals. Elife 10 (2021).

17. J. Y. Lu, M. B. Sewer, p54nrb/NONO regulates cyclic AMP-dependent glucocorticoid production by modulating phosphodiesterase mRNA splicing and degradation. Mol Cell Biol 35, 1223–1237 (2015).

18. J. C. Schwartz et al., FUS binds the CTD of RNA polymerase II and regulates its phosphorylation at Ser2. Genes Dev 26, 2690–2695 (2012).

19. M. Mikula, K. Bomsztyk, K. Goryca, K. Chojnowski, J. Ostrowski, Heterogeneous nuclear ribonucleoprotein (HnRNP) K genome-wide binding survey reveals its role in regulating 3’-end RNA processing and transcription termination at the early growth response 1 (EGR1) gene through XRN2 exonuclease. J Biol Chem 288, 24788–24798 (2013).

20. T. Hirose et al., NEAT1 long noncoding RNA regulates transcription via protein sequestration within subnuclear bodies. Mol Biol Cell 25, 169–183 (2014).

21. M. Modic et al., Cross-Regulation between TDP-43 and Paraspeckles Promotes Pluripotency-Differentiation Transition. Mol Cell 74, 951–965 e913 (2019).

22. D. Reddy et al., Paraspeckles interact with SWI/SNF subunit ARID1B to regulate transcription and splicing. EMBO Rep 24, e55345 (2023).

23. C. Wang et al., Stress Induces Dynamic, Cytotoxicity-Antagonizing TDP-43 Nuclear Bodies via Paraspeckle LncRNA NEAT1-Mediated Liquid-Liquid Phase Separation. Mol Cell 79, 443–458 e447 (2020).

24. P. Zhang, L. Cao, R. Zhou, X. Yang, M. Wu, The lncRNA Neat1 promotes activation of inflammasomes in macrophages. Nat Commun 10, 1495 (2019).

25. H. Ma et al., The Long Noncoding RNA NEAT1 Exerts Antihantaviral Effects by Acting as Positive Feedback for RIG-I Signaling. J Virol 91 (2017).

26. Q. Zhang, C. Y. Chen, V. S. Yedavalli, K. T. Jeang, NEAT1 long noncoding RNA and paraspeckle bodies modulate HIV-1 posttranscriptional expression. mBio 4, e00596–00512 (2013).

27. F. Prinz, A. Kapeller, M. Pichler, C. Klec, The Implications of the Long Non-Coding RNA NEAT1 in Non-Cancerous Diseases. Int J Mol Sci 20 (2019).

28. K. Imamura et al., Long Noncoding RNA NEAT1-Dependent SFPQ Relocation from Promoter Region to Paraspeckle Mediates IL8 Expression upon Immune Stimuli. Mol Cell 54, 1055 (2014).

29. J. Saini, U. Thapa, B. Bandyopadhyay, S. Vrati, A. Banerjee, Knockdown of NEAT1 restricts dengue virus replication by augmenting interferon alpha-inducible protein 27 via the RIG-I pathway. J Gen Virol 104 (2023).

30. T. Naganuma, T. Hirose, Paraspeckle formation during the biogenesis of long non-coding RNAs. RNA Biol 10, 456–461 (2013).

31. Y. S. Mao, H. Sunwoo, B. Zhang, D. L. Spector, Direct visualization of the co-transcriptional assembly of a nuclear body by noncoding RNAs. Nat Cell Biol 13, 95–101 (2011).

32. H. An, J. T. Tan, T. A. Shelkovnikova, Stress granules regulate stress-induced paraspeckle assembly. J Cell Biol 218, 4127–4140 (2019).

33. K. Imamura et al., Long noncoding RNA NEAT1-dependent SFPQ relocation from promoter region to paraspeckle mediates IL8 expression upon immune stimuli. Mol Cell 53, 393–406 (2014).

34. P. J. Murray, Macrophage Polarization. Annu Rev Physiol 79, 541–566 (2017).

35. T. Hirose, T. Yamazaki, S. Nakagawa, Molecular anatomy of the architectural NEAT1 noncoding RNA: The domains, interactors, and biogenesis pathway required to build phase-separated nuclear paraspeckles. Wiley Interdiscip Rev RNA 10, e1545 (2019).

36. V. R. Ramirez-Carrozzi et al., Selective and antagonistic functions of SWI/SNF and Mi-2beta nucleosome remodeling complexes during an inflammatory response. Genes Dev 20, 282–296 (2006).

37. T. Kawaguchi et al., SWI/SNF chromatin-remodeling complexes function in noncoding RNA-dependent assembly of nuclear bodies. Proc Natl Acad Sci U S A 112, 4304–4309 (2015).

38. K. O. West et al., The Splicing Factor hnRNP M Is a Critical Regulator of Innate Immune Gene Expression in Macrophages. Cell Rep 29, 1594–1609 e1595 (2019).

39. M. B. Clark et al., Genome-wide analysis of long noncoding RNA stability. Genome Res 22, 885–898 (2012).

40. D. Strassheim et al., Phosphoinositide 3-kinase and Akt occupy central roles in inflammatory responses of Toll-like receptor 2-stimulated neutrophils. J Immunol 172, 5727–5733 (2004).

41. T. Kawai, S. Akira, Signaling to NF-kappaB by Toll-like receptors. Trends Mol Med 13, 460–469 (2007).

42. B. E. Aubol et al., N-terminus of the protein kinase CLK1 induces SR protein hyperphosphorylation. Biochem J 462, 143–152 (2014).

43. J. Prasad, J. L. Manley, Regulation and substrate specificity of the SR protein kinase Clk/Sty. Mol Cell Biol 23, 4139–4149 (2003).

44. J. Houseley, J. LaCava, D. Tollervey, RNA-quality control by the exosome. Nat Rev Mol Cell Biol 7, 529–539 (2006).

45. T. Tanu et al., hnRNPH1-MTR4 complex-mediated regulation of NEAT1v2 stability is critical for IL8 expression. RNA Biol 18, 537–547 (2021).

46. S. Nakagawa, T. Naganuma, G. Shioi, T. Hirose, Paraspeckles are subpopulation-specific nuclear bodies that are not essential in mice. J Cell Biol 193, 31–39 (2011).

47. S. Nakagawa et al., The lncRNA Neat1 is required for corpus luteum formation and the establishment of pregnancy in a subpopulation of mice. Development 141, 4618–4627 (2014).

48. R. Elling et al., Genetic Models Reveal cis and trans Immune-Regulatory Activities for lincRNA-Cox2. Cell Rep 25, 1511–1524 e1516 (2018).

49. N. A. Eisele et al., Salmonella require the fatty acid regulator PPARdelta for the establishment of a metabolic environment essential for long-term persistence. Cell Host Microbe 14, 171–182 (2013).

50. S. J. Taylor, S. E. Winter, Salmonella finds a way: Metabolic versatility of Salmonella enterica serovar Typhimurium in diverse host environments. PLoS Pathog 16, e1008540 (2020).

51. S. K. Lathrop et al., Replication of Salmonella enterica Serovar Typhimurium in Human Monocyte-Derived Macrophages. Infect Immun 83, 2661–2671 (2015).

52. A. Vazquez-Torres, J. Jones-Carson, P. Mastroeni, H. Ischiropoulos, F. C. Fang, Antimicrobial actions of the NADPH phagocyte oxidase and inducible nitric oxide synthase in experimental salmonellosis. I. Effects on microbial killing by activated peritoneal macrophages in vitro. J Exp Med 192, 227–236 (2000).

53. A. Vazquez-Torres et al., Toll-like receptor 4 dependence of innate and adaptive immunity to Salmonella: importance of the Kupffer cell network. J Immunol 172, 6202–6208 (2004).

54. M. Modolell, I. M. Corraliza, F. Link, G. Soler, K. Eichmann, Reciprocal regulation of the nitric oxide synthase/arginase balance in mouse bone marrow-derived macrophages by TH1 and TH2 cytokines. Eur J Immunol 25, 1101–1104 (1995).

55. C. I. Chang, B. Zoghi, J. C. Liao, L. Kuo, The involvement of tyrosine kinases, cyclic AMP/protein kinase A, and p38 mitogen-activated protein kinase in IL-13-mediated arginase I induction in macrophages: its implications in IL-13-inhibited nitric oxide production. J Immunol 165, 2134–2141 (2000).

56. A. J. Tong et al., A Stringent Systems Approach Uncovers Gene-Specific Mechanisms Regulating Inflammation. Cell 165, 165–179 (2016).

57. K. Imamura et al., Diminished nuclear RNA decay upon Salmonella infection upregulates antibacterial noncoding RNAs. EMBO J 37 (2018).

58. L. Davidson et al., Rapid Depletion of DIS3, EXOSC10, or XRN2 Reveals the Immediate Impact of Exoribonucleolysis on Nuclear RNA Metabolism and Transcriptional Control. Cell Rep 26, 2779–2791 e2775 (2019).

